# Leaf area index estimation of even-aged oak (*Quercus petraea*) forests using in situ stand dendrometric parameters

**DOI:** 10.1101/2021.08.05.454476

**Authors:** M. Briere, C. François, F. Lebourgeois, I. Seynave, G. Vincent, N. Korboulewsky, F. Ningre, T. Perot, S. Perret, A. Calas, E. Dufrêne

## Abstract

The leaf area index (LAI) is a key characteristic of forest stand aboveground net productivity (ANP), and many methods have been developed to estimate the LAI. However, every method has flaws, e.g., methods may be destructive, require means or time and/or show intrinsic bias and estimation errors.

A relationship using basal area (G) and stand age to estimate LAI was proposed by Sonohat et al. (2004). We used literature data in addition to data form measurements campaign made in the northern half of France to build a data set with large ranges of pedoclimatic conditions, stand age and measured LAI. We validated the Sonohat et al. (2004) relationship and attempted to improve or modify it using other stand/dendrometric characteristics that could be predictors of the LAI.

The result is a series of three models using the G, age and/or quadratic mean diameter (Dg), and the models were able to estimate the LAI of an oak only even-aged forest stand with good confidence (root mean square error, RMSE < 0.75) While G is the main predictor here, age and Dg could be used conjointly or exclusively given the available data, with variable precision in the estimations.

Although these models could not, by construction, relate to the interannual variability of the LAI, they may provide the theoretical LAI of an untouched forest (no meteorological, biotic or anthropogenic perturbation) in recent years. additionally, the use of this model may be more interesting than an LAI measurement campaign, depending on the means to be invested in such a campaign.

## Introduction

Forest productivity in the near future is a subject of great concern (Lindner et al. 2010; Hanewinkel et al. 2013; Hurlbert et al. 2019), and the identification of functional traits, or their combination, connected to tree adaptation to climate change is still under debate (Bussotti et al. 2015). Aboveground net productivity (ANP) is driven by many intrinsic (e.g., light-use efficiency, drought resistance, leaf area, species composition) and extrinsic (e.g., nutrient availability, meteorology, topography, management) factors (Skovsgaard and Vanclay 2008). In this context, stand characteristics are important variables to model ANP (Stage 1973; Peng 2000; Monserud 2003); canopy description is especially important, as it is the main interface between a tree and the atmosphere. Its description has been subject to research, and several descriptors have been proposed (Jonckheere et al. 2004; Parker 2020),such as the leaf area index (LAI) (Running and Coughlan 1988; Asner et al. 2003; Clark et al. 2008; Ollinger 2011; Parker 2020). The LAI is by far the main variable used to describe a canopy, and is one of the main factors influencing the ANP of forest stands (Fassnacht and Gower 1997; Barr et al. 2004; Reich 2012).

The LAI is defined as half the total leaf area developed by the vegetation reported on the land surface occupied by this vegetation (Chen and Black 1992). The LAI is an important value used to estimate the water and radiative budget of a forest stand and thus stand productivity (Running and Coughlan 1988; Landsberg and Waring 1997; Pietsch et al. 2005). The LAI has a great impact on forests since higher LAI values indicates greater transpiration and greater rain interception than lower LAI values (Bréda and Granier 1996; Granier et al. 1999; Yan et al. 2012). Considering the radiative budget, a lower LAI indicates less sunlight absorption, which would have two direct effects: (i) lower photosynthetic activity leading to decreases in tree productivity and (ii) higher production of understorey vegetation (Granier et al. 1999), leading to the development of understorey vegetation, often accompanied by higher evapotranspiration and competition for soil water (Black et al. 1989). Quantitatively speaking, some studies have proposed linear relationships between ANP and LAI with linear factors of 0.3 to 1.9 Mg·ha^-1^·year^-1^ per LAI unit (Fassnacht and Gower 1997; Asner et al. 2003; Parker 2020) or a logistic relationship (Jokela et al. 2004).

Nevertheless, LAI measurements require the implementation of heavy protocols. LAI measurement methods are divided into two main categories: (i) direct methods, implying a measurement of leaf area, which can be used to define an allometric relationship to extend the measure made on a subsample to the whole sample (in which case, the term semidirect method is preferred) and (ii) indirect methods, which consist of the analysis of the light under the canopy with the assumption of spatial and angular distribution of leaves (Dufrêne and Bréda 1995; Bréda 2003; Jonckheere et al. 2004; Weiss et al. 2004).

Among direct methods, **litter traps** are often considered as reference (Dufrêne and Bréda 1995; Mussche et al. 2001; Bréda et al. 2002; Bréda 2003). This method consists of the placement, of mesh-bottomed (or, at least, draining material) collectors of determined size, shape and height just above the ground, and these traps collect leaves during the leaf fall season (Morrison 2011). This method requires the installation of enough litter traps and regular collection during the leaf fall season; additionally, samples require sorting, weighting and measuring the leaf surface of the litter collected to avoid the loss of material by decomposition or degradation (Bréda et al. 2002; Bréda 2003; Eriksson et al. 2005). A common semidirect method based on **allometry** consists of the construction of a relationship between easily measurable tree characteristics such as diameter at breast height or stem basal area and the leaf surface of the tree (Bartelink 1997; Le Dantec et al. 2000). However, this method requires cutting down the selected trees and collecting and measuring all of the tree’s leaves. The relation between the measured variables and leaf area is then applied to the other trees of the stand, and the total leaf area is estimated. A third method, **inclined point quadrats**, is applicable only for herbaceous stands and consists of planting a thin probe regularly in the leaf coverage and counting the contacts between the probe and the leaves (Wilson 1960).

Finally, the **needle method** is derived from the inclined point quadrats method applied for deciduous trees at the end of fall when all leaves are on the ground. Under these conditions all leaves are horizontal and flat, and the probe is vertical. In this case, the number of contact points between the probe and the leaves is locally equal to the LAI. The mean value of a large number of measurements in a forest stand gives the mean LAI of the stand (Bréda et al. 2002; Bréda 2003).

Indirect methods use light as a tool to estimate the LAI and rely on two theories: gap fraction and gap size distribution (Chen and Cihlar 1995). The simplest application of these theories is to measure sun flecks on the ground multiple times a day. The proportion of sunlit ground over time (gap fraction) or the distribution of the sizes of this fleck (gap-size distribution) can provide enough information to estimate the LAI (Welles 1990; Welles and Cohen 1996). An evolution of this method is the d**igital hemispherical photography (DHP)**. With a shaded canopy and a homogeneously lit the sky (during dawn or dusk or a cloudy day), it is possible to distinguish sky (lightened areas) from the leaves (dark areas). Using gap fraction or gap-size distribution theories, it is possible to estimate the LAI (Weiss et al. 2004). Many instruments use these theories to indirectly measure the LAI under different types of canopies (from prairies to forests), such as the **Licor LAI-2000**, the Demon, the Plant Canopy Imager, or a smartphone with the “Pocket LAI” application (Bréda et al. 2002; Jonckheere et al. 2004; Casa et al. 2019).

Finally, other methods based solely on modelling allow the estimation of the LAI (Running and Gower 1991; Pietsch et al. 2005; Guillemot et al. 2017). This approach allows the models to estimate the interannual variation in the LAI and, thus, the variation in the energy, water and carbon budget of the stand. Application to a real stand would require some variable that is easily accessible using a model but harder to access in a real-life forest stand (i.e., net assimilation of carbon or quantity of available soil water).

None of these techniques are perfect and each has flaws. While litter trapping and allometry can distinguish the specific LAI of different tree species in the canopy, indirect methods cannot (Bréda 2003). For an LAI higher than 5, instruments using gap fraction theory show a saturation in the measurements (Gower et al. 1996). Some direct or semidirect methods require either destructive measurements of trees or the installation of materials on the stand (litter traps, sensors and data acquiring unit) while others require a large amount of time spent on the stand to achieve a correct sampling. This large amount of recquiered effort and/or time may lead to subsampling and, thus, a biased estimation of the measured LAI. Last, assumptions are made for each of these methods, especially for indirect ones, and they must be considered when considering the results because these assumptions are often not verifiable.

LAI values are difficult to estimate directly, and it appears important, as an alternative to measurements, to develop a relationship linking the LAI to common stand parameters through simple empirical relationships. Sonohat et al. (2004) developed such a relationship to estimate the LAI of a coniferous stand using the assumption of an age-dependent relationship between the basal area (G) and LAI. Based on this study, we propose several adaptations of tis relationship to improve the LAI estimation for even-aged oak stands using dendrometric stand data: G, stand age and/or quadratic mean diameter (Dg). To establish and validate our relationships, different data sets e compiled, including an LAI measuring campaign, using the needles method, through northern France during the winters of 2018-2019 and 2019-2020 (35 plots distributed on 13 sites); this campaign was supplemented by 86 published measures (distributed on 44 sites) from the literature (Le Dantec et al. 2000; Balandier et al. 2006) and 22 measures (distributed on 4 sites) from personal correspondence, resulting in a total of 145 measures (See Appendix 1).

## Material and Methods

### LAI measurement and estimation

#### Summary of the data sets

A total of 145 data points were considered in our study and came from different datasets. This compiled dataset covered a wide range of densities (Relative Density Index (relative density index (RDI) (Reineke 1933; Le Goff et al. 2011; Le Moguédec and Dhôte 2012)): 0.18 to 1.37; mean = 0.65) and ages (10 to 220; mean = 110 years). This thorough approach is quite an achievement since only a few LAI measurement data are available on pure *Q. petraea* stands and fewer data sets contain stand-related variables such as G or tree densities. The ranges [min-max (mean)] of G, Dg and LAI were 6.9-49 (21.6) m^2^/ha, 2.8-80.9 (36.1) cm and 0.4-5.6 (3.1) m^2^/m^2^, respectively (see Appendix 1).

The LAI was measured by either a direct or an indirect method: litter-trap followed by leaf surface measurements, needle method, DHP, transmittance and LAI-2000 (see Appendix 1).

#### Description of the needle method

The needle method is derived from the inclined quadrat points (Wilson 1960). This derivation implies flat horizontal leaves and a vertical thin probe. For this measurement we used a square-shaped (∼10×10cm) plastic plate pierced by a hole (diameter = 0.8 mm) on its centre and a thin (diameter = 0.8 mm, 20 cm) long steel needle (piano string) as the probe. For the measurement of each plot, at least two transects (usually diagonals) were planned. The distance between throws was estimated on site to cover the length of the whole transects and we obtained between 117 and 163 (mean = 134) local LAI measurements.

Along this transect, plastic plates were thrown to the soil (backwards to ensure minimization of the operator effect on the measurement). For each throw, the plate was pushed flat, and litter was stabbed through the plate’s centre hole. The plates were then carefully picked up and flipped over. As stated by (Parker 2020) “In the idealized concept, LAI is the number of leaves above a single point, that contact an infinitely thin vertical line”. Each contact point, defined by the occurrence of the probe passing through or touching the border of a leaf, is counted, identified to the leaf species and recorded down. The main limitations of this technique are the shape of the leaf and the diameter of the probe. A large probe diameter and/or a leaf showing a high perimeter over area ratio might increase the border effect and may bias the measurement. The diameter of the probe (0.8 mm diameter stainless steel piano chord) and the shape of the oak leaves minimize bias. The LAI of the stand is computed, and then the mean value of all stabs can be discriminated by species. For practical reasons, the two most frequent species in each stand were distinguished, and other species were pooled into the “other category”. The data resulting from this measurement campaign are detailed in Table 1 (details available in Appendix 2). Over this measurement campaign, an average of 139 local LAI measurement were done each of the 35 experimental plots (total = 4809, mean value = 3, mean error on measurement = 0.4) (more details are supplied in the Appendix 2).

**Table 1:** Summary of LAI measures using the needles technique. The LAI is the mean value of all measurements on each plot.

| Network | Site | plot | Latitude | Longitude | RDI | Age<sub>2018</sub><br>(years) | Density<br>(N. ha<sup>-1</sup>) | G<br>(m<sup>2</sup>. ha<sup>-1</sup>) | Dg<br>(cm) | LAI<br>(m<sup>2</sup>. m<sup>-2</sup>) |
| --- | --- | --- | --- | --- | --- | --- | --- | --- | --- | --- |
| ICOS | Barbeau |  | 48.4764 | 2.7801 | 0.56 | 134 | 119 | 22.2 | 38.1 | 3.1 |
| GIS Coop | Darney-Sessile | 1 | 48.0954 | 6.1316 | 0.22 | 32 | 168 | 7.7 | 23.9 | 1.8 |
| GIS Coop | Darney-Sessile | 2 | 48.0954 | 6.1316 | 0.89 | 32 | 1627 | 25.7 | 14.5 | 2.4 |
| GIS Coop | Darney-Sessile | 3 | 48.0954 | 6.1316 | 0.87 | 33 | 1585 | 24.6 | 14.4 | 2.9 |
| GIS Coop | Darney-Sessile | 4 | 48.0954 | 6.1316 | 0.42 | 33 | 548 | 13.5 | 17.6 | 2.5 |
| GIS Coop | Grosbois | 1 | 46.4999 | 2.9930 | 0.47 | 37 | 656 | 14.9 | 16.9 | 3.5 |
| GIS Coop | Montrichard | 2 | 47.3751 | 1.1784 | 1.33 | 33 | 3065 | 25.7 | 10.3 | 3.5 |
| GIS Coop | Montrichard | 4 | 47.3751 | 1.1784 | 0.50 | 35 | 916 | 14.9 | 14.4 | 2.7 |
| GIS Coop | Montrichard | 5 | 47.3751 | 1.1784 | 0.85 | 32 | 2470 | 22.4 | 10.6 | 2.7 |
| GIS Coop | Moulins-Bonsmoulins | 1 | 48.6956 | 0.5187 | 0.64 | 25 | 974 | 12.8 | 12.8 | 3.2 |
| GIS Coop | Parroy | 1 | 48.6702 | 6.6523 | 1.11 | 41 | 1451 | 29.9 | 17.6 | 3.7 |
| GIS Coop | Parroy | 2 | 48.6702 | 6.6523 | 0.85 | 40 | 963 | 25.3 | 19.1 | 3.1 |
| GIS Coop | Parroy | 3 | 48.6702 | 6.6523 | 0.35 | 40 | 120 | 11.3 | 38.5 | 3.4 |
| GIS Coop | Parroy | 4 | 48.6702 | 6.6523 | 0.22 | 39 | 100 | 10.0 | 32.3 | 3.0 |
| GIS Coop | Reno-Valdieu | 1 | 48.5276 | 0.6780 | 0.51 | 52 | 339 | 18.2 | 26.3 | 2.7 |
| GIS Coop | Reno-Valdieu | 2 | 48.5276 | 0.6780 | 1.27 | 52 | 1389 | 33.4 | 19.5 | 2.9 |
| GIS Coop | Reno-Valdieu | 3 | 48.5276 | 0.6780 | 0.70 | 51 | 669 | 22.3 | 21.2 | 2.9 |
| GIS Coop | Reno-Valdieu | 4 | 48.5276 | 0.6780 | 0.25 | 52 | 120 | 10.1 | 31.8 | 2.5 |
| GIS Coop | Tronçais | 1 | 46.6636 | 2.7072 | 0.45 | 56 | 333 | 16.3 | 24.5 | 3.2 |
| GIS Coop | Tronçais | 2 | 46.6636 | 2.7072 | 0.24 | 57 | 129 | 9.1 | 29.6 | 3.0 |
| GIS Coop | Tronçais | 3 | 46.6636 | 2.7072 | 1.04 | 55 | 1345 | 30.6 | 17.7 | 3.6 |
| OPTMix | O12 | 2 | 47.7737 | 2.5755 | 0.37 | 76 | 294 | 12.8 | 23.6 | 3.4 |
| OPTMix | O12 | 3 | 47.7737 | 2.5755 | 0.62 | 76 | 500 | 21.5 | 23.4 | 3.8 |
| OPTMix | O214 | 1 | 47.8081 | 2.4468 | 0.60 | 68 | 604 | 19.9 | 20.5 | 3.9 |
| OPTMix | O214 | 2 | 47.8081 | 2.4468 | 0.37 | 68 | 280 | 12.8 | 24.1 | 3.3 |
| OPTMix | O593 | 1 | 47.9127 | 2.3012 | 0.62 | 67 | 425 | 21.9 | 25.6 | 4.0 |
| OPTMix | O593 | 2 | 47.9127 | 2.3012 | 0.34 | 67 | 192 | 12.6 | 28.9 | 3.6 |
| LERFoB | Blois (Sablonnières) | 1 | 47.5705 | 1.2573 | 0.37 | 129 | 102 | 16.1 | 49.1 | 2.7 |
| LERFoB | Blois (Sablonnières) | 2A | 47.5705 | 1.2573 | 0.24 | 129 | 104 | 9.3 | 49.2 | 2.3 |
| LERFoB | Blois (Sablonnières) | 2B | 47.5705 | 1.2573 | 0.52 | 129 | 377 | 18.1 | 36.1 | 2.5 |
| LERFoB | Blois (Sablonnières) | 3 | 47.5705 | 1.2573 | 0.86 | 129 | 278 | 34.7 | 39.9 | 2.5 |
| LERFoB | Blois (Sablonnières) | 4 | 47.5705 | 1.2573 | 0.65 | 129 | 184 | 27.1 | 43.3 | 2.5 |
| LERFoB | Tronçais (Tresor) | 1 | 46.6703 | 2.7224 | 1.02 | 139 | 329 | 41.4 | 40.0 | 3.4 |
| LERFoB | Tronçais (Tresor) | 2 | 46.6703 | 2.7224 | 0.85 | 139 | 279 | 34.2 | 39.5 | 3.0 |
| LERFoB | Tronçais (Tresor) | 3 | 46.6703 | 2.7224 | 0.62 | 139 | 152 | 26.3 | 47.0 | 2.8 |

To assess the required sampling effort and the performance of the needle method, we performed a study based on a representative distribution of leaf contact points. For this purpose, we assumed that the actual measurements made along the transects were representative of the whole plot. Then, a distribution function was fitted to match the distribution of measurements (Figure 1).

**Figure 1:**
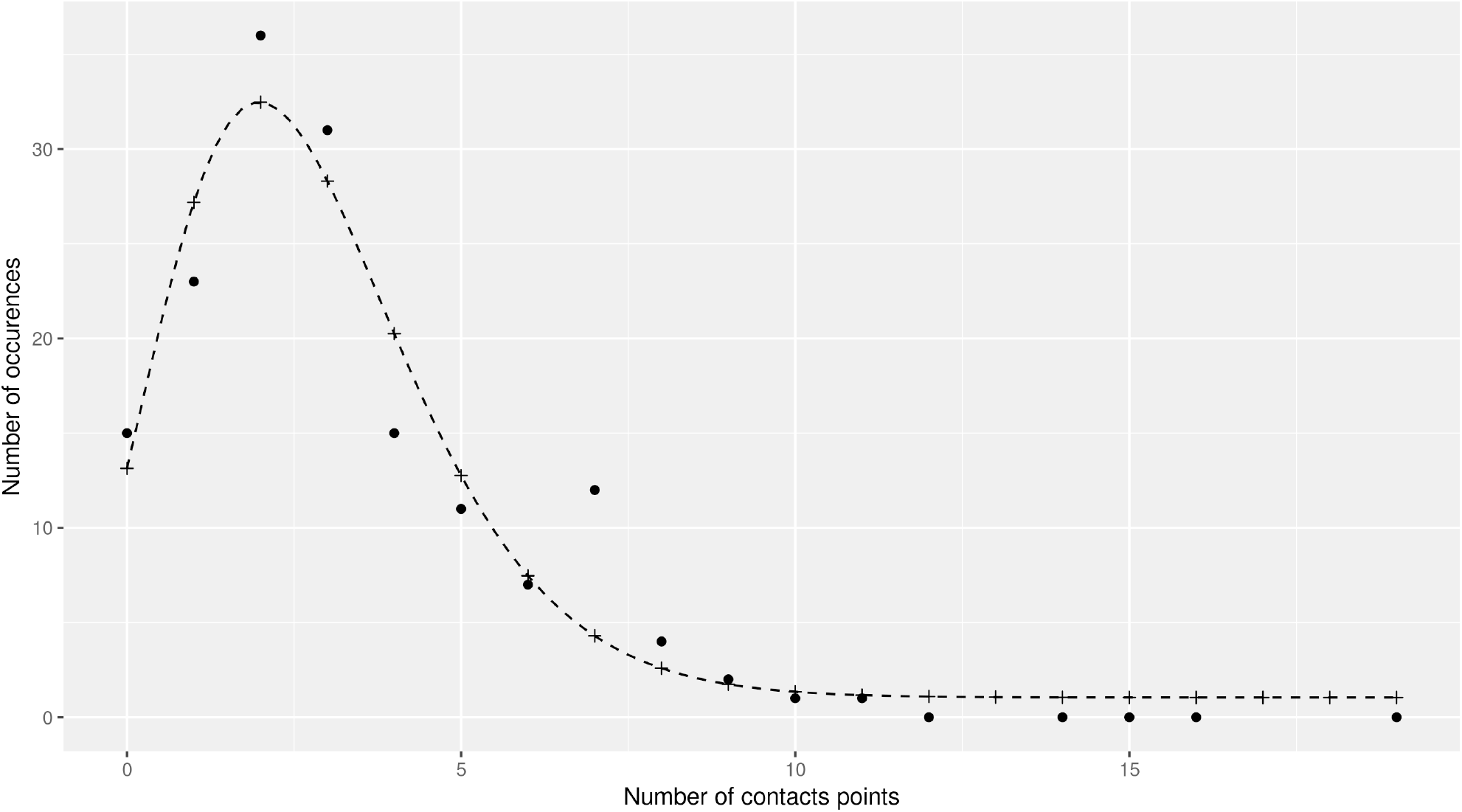
Occurrences of the local LAI values observed. Points are the field data while crosses represents the values obtained by the application of the function fitted.

This distribution has been used to generate a virtual sample of ten thousand measurements, and several sampling schemes have been studied. For each simulated sampling effort (from n=1 to n=500), a thousand random draws (with replacement) were generated in this virtual sample to evaluate the mean dispersion given the number of throws (sample size) made.

#### Determination of oak-only LAI from stand LAI when indirect methods are used

One of the main drawbacks of indirect methods is their inability to distinguish between species when measuring the LAI. In their study, (Genet et al. 2010) proposed a model to estimate the LAI belonging to oaks in a mixed stand (*Q. pertaea* and *Fagus sylvatica L*.) (Eq. (1)).

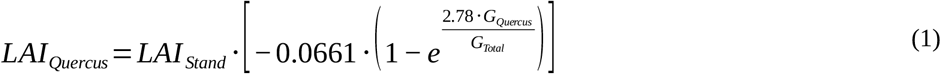

where LAI_Quercus_ and LAI_Stand_ are the LAIs of oaks only and of the whole stand, respectively, and G_Quercus_ and G_Total_ are the basal areas of oaks only and of the whole stand, respectively.

#### Computation of LAI from measured transmittance

It appears that a subset of our gathered database was comprised of below-canopy transmittance measurements (instead of LAI measurements). To convert these transmittances into LAI estimations we used a modelization and look-up table (LUT) approach. The estimation of the transmittance by the canopy can be approached using the Beer-Lambert relationship with an extinction coefficient relative to the LAI (Eq. (2)). The extinction coefficient includes leaf optical properties, leaf angle distribution and a clumping index (Vose et al. 1995; Cannell and Grace 2011). To estimate this extinction coefficient, we used the scattering by arbitrarily inclined leaves (SAIL) model (Verhoef 1984) in a multilayer version as described in Dufrêne et al. (2005). In this implementation, the canopy is divided into a stack of discrete layers. Each layer has a depth of 0.2 LAI unit (a forest with LAI=6 therefore has 30 layers). In our version of the SAIL model, the woody parts of the canopy are estimated to represent 0.9 LAI units (Dufrêne and Bréda 1995; Cutini et al. 1998). The radiation extinction and diffusion of each single layer (elementary reflectances and transmittances) are based on the SAIL model (Verhoef 1984, 1985).

The transmittances were then computed for a whole range of LAI values (1.1 - 8) under specific conditions (80% diffuse radiation, day 185, average between 10 am and 4 pm, according to the measurement conditions described in Balandier et al., 2006. An extinction coefficient (k) was then fitted using the Beer-Lambert law to obtain a relationship between transmittance and LAI:

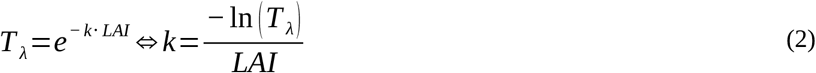

This relationship allows for the transformation of measured transmittance values into LAI estimations through a (LUT) between transmittances and LAI.

### Experimental design and data collecting

A campaign of LAI measurements using the needle method was carried out during the winters of 2018-2019 and 2019-2020 at 13 even-aged sessile oak (*Quercus petraea (Matt.) Liebl*.) experimental sites distributed over the northern half of France (see circles in Figure 2 and Table 1, supplementary data in Appendix 2). Each site was composed of plots with different stand density treatments: overall, 35 plots were measured.

**Figure 2:**
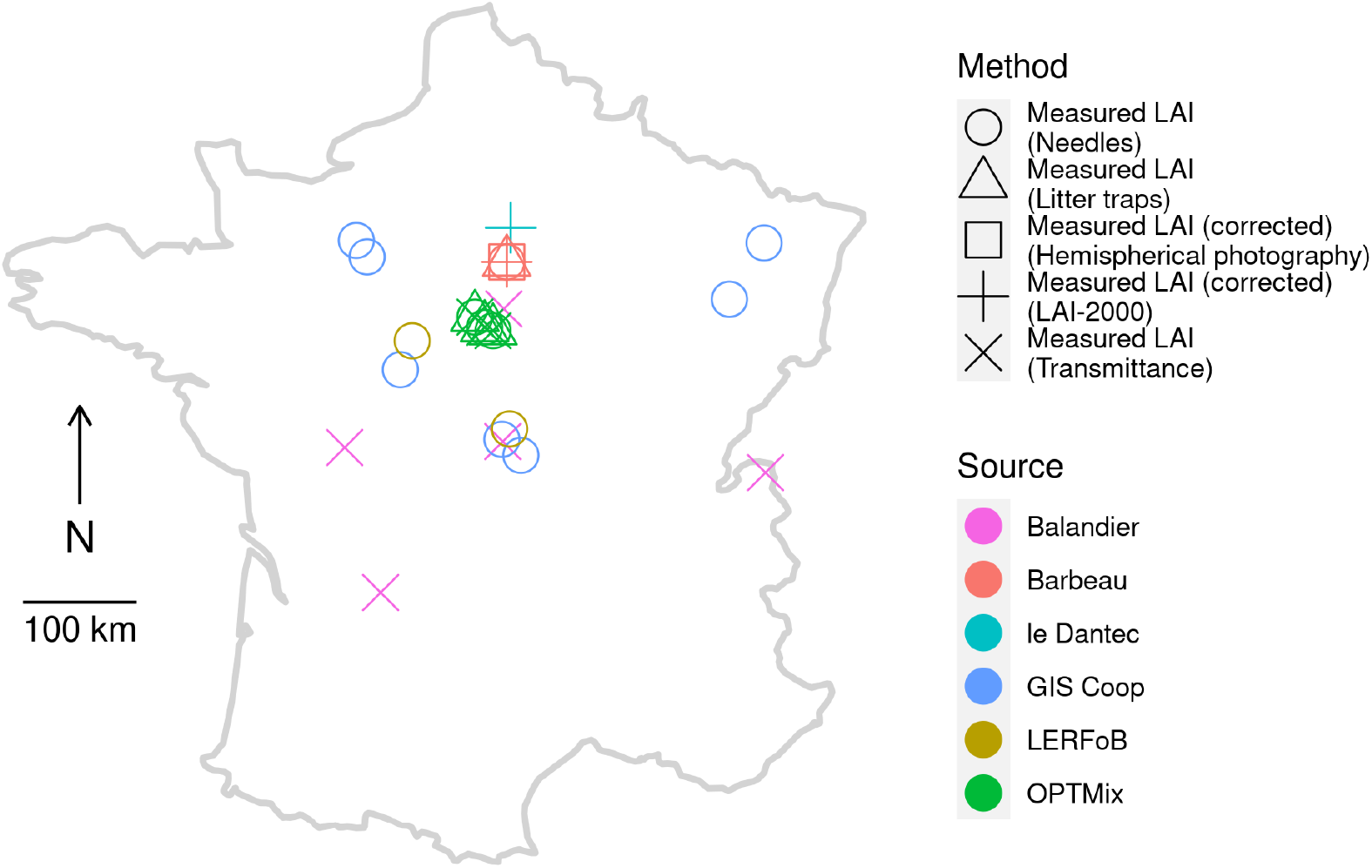
Locations of LAI measurements summarized in Table 1. Colors represents either the origin of the data or the network managing the experimental plots on which needles measurements have been made. The shape represents the method used to measure the LAI on each plot.

**Figure 3:**
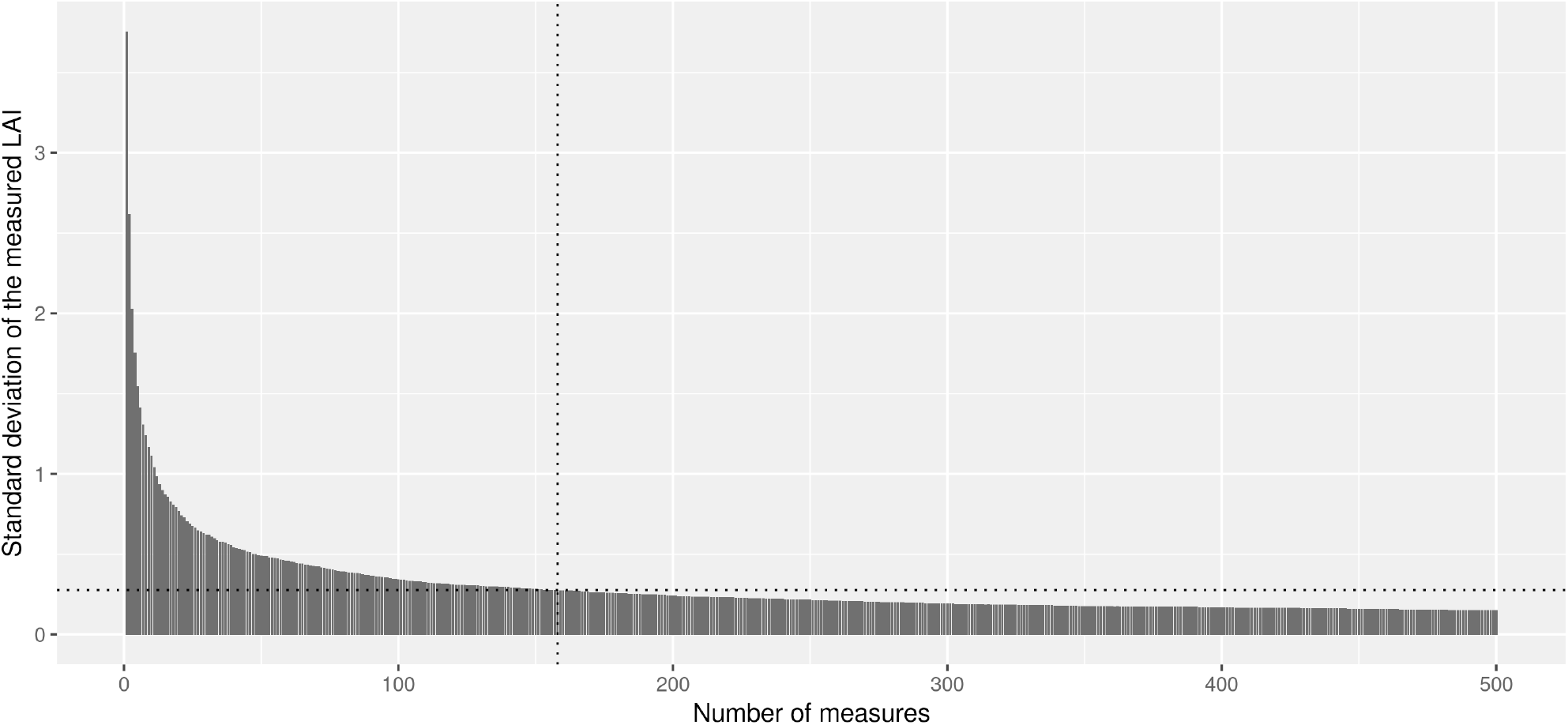
Evolution of the uncertainty of the measure given the number of throws made on the plot. The dotted lines mark the number of measures to achieve to obtain a twice as much precision.

The needle method was used on all sites labelled here as LERFoB, GISCoop, OPTMix and Barbeau. The method allowed us to count the leaves according to their species and therefore to estimate the oak’s LAI, excluding other species. The characteristics of each plot are presented in Table 1.

#### Experimental sites 1: LERFoB network

The LERFoB oak-permanent-plot network, which consists of thinning trials, aimed at analysing the effects of competition on tree growth and stand production in even-aged, naturally regenerated sessile oak (*Q. petraea*), stands across northern France (https://www6.inrae.fr/in-sylva-france/Services/In-Situ/Reseau-LERFOB)

It was installed between 1925 and 1956, in five forest areas, in mostly pure even-aged stands, on soils favourable to oak growth; these factors could be assessed by the dominant height, which corresponded to a medium to good site index (Duplat and Tran-Ha 1997). This network covers a significant part of the sessile oak distribution in continental France.

This network of long-term experiments, which is still being monitored, covers a wide range of silvicultural treatments and presented contrasted initial ages, between 27 and 121 years (Oswald 1981). Different thinning intensities were compared in each trial.

The sampled plots were chosen at two sites where the largest differences, in RDI value, were observed: Sablonnières (Blois State Forest) and Trésor (Tronçais State Forest) (Table 1, supplementary data in Appendix 1).

#### Experimental sites 2: GIS Coop

The GIS Coop network has a wide geographical spread across the northern half of France (Sindou et al 2001; Seynave et al. 2018). For each site, at least three experimental plots were present with different stand densities: from free growth (RDI_goal_ < 0.2) to self-thinning densities (RDI_goal_ > 0.9) with intermediate values (RDI_goal_ ≃ 0.25 and RDI_goal_ ≃ 0.5) and variable RDI_goal_ (ascending or descending values with time). This experimental design provided a wide range of basal areas (from 7.7 to 33.4 m^2^.ha^-1^), with particularly very low values, and allowed us to decorrelate stand age and stand dendrometric data (Dg, G and N). The plots were from 25 to 57 years old at the time of measurements (Table 1, supplementary data in Appendix 1).

The presence and density of the understorey varied with site and treatment. Low-density treatments often showed higher understorey density than high-density treatments. This understorey is regularly controlled to limit its development.

#### Experimental sites 3: OPTMix

The three OPTMix sites were all located in the Orléans state forest and presented each two density treatments with objective RDI_goal_ ≃ 0.4 and ≃ 0.6 (Table 1). The understorey was absent and every species other than *Q. petraea* was removed. In addition to our measures using the needle method, LAI measurements were made on OPTMix plots for the 2018 vegetation season year using litter traps and global radiation transmittance (Table 2) (Korboulewsky et al. 2015; Perot et al. 2019)

**Table 2:** Data from the OPTMix network. All values necessary for estimating the LAI using our model, (G, age, Dg), plus additional information (Density), are given. The columns marked with a star (*) are calculated values based on measurements. The LAI “transmittance” is the result of the transformation of total solar radiation transmittance (T_TSR_) measurement to LAI using the Beer-Lambert relationship detailed below (see Eq (2)).

| Id | RDI | Age | Density<br>($n \cdot ha^{-1}$) | G<br>($m^2 \cdot ha^{-1}$) | Dg*<br>(cm) | $T_{TSR}$<br>(%) | $LAI_{quercus}$<br>($m^2 \cdot m^{-2}$) | | |
| --- | --- | --- | --- | --- | --- | --- | --- | --- | --- |
|  |  |  |  |  |  |  | Litter traps | Transmittance* | Needles |
| O12-2 | 0.35 | 76 | 294 | 12.8 | 23.6 | 23% | 2.9 | 3.0 | 3.4 |
| O12-3 | 0.59 | 76 | 500 | 21.5 | 23.4 | 9% | 3.7 | 4.8 | 3.8 |
| O214-1 | 0.53 | 68 | 604 | 19.9 | 20.5 | 10% | 3.7 | 4.5 | 3.9 |
| O214-2 | 0.35 | 68 | 280 | 12.8 | 24.1 | 23% | 2.7 | 2.9 | 3.3 |
| O593-1 | 0.60 | 67 | 425 | 21.9 | 25.6 | 15% | 3.9 | 3.7 | 4.0 |
| O593-2 | 0.35 | 67 | 192 | 12.6 | 28.9 | 44% | 2.8 | 1.6 | 3.6 |

#### Experimental site 4: Barbeau

The Barbeau forest hosts an experimental site of the Integrated Carbon Observation System (ICOS, https://www.icos-cp.eu) (Delpierre et al. 2016). The vegetation is composed of mature oaks of age 135 years and hornbeams (*Carpinus betulus*) as the understorey. Although there is an understorey, the basal areas and LAI were measured separately for each of these species (using litter traps). For this study, only the oak part of the LAI measurements was considered. Other LAI measurements were available for this plot, Li-cor LAI-2000 and DHP (Table 3) (Montagnani 2018). Because these measures were unable to separate LAIs from different species, the calculations were made using the model proposed by Genet et al. (2010) (Eq. (1)).

**Table 3:** Data from the ICOS site of Barbeau. All values necessary for estimating the LAI using our model, (G, age, Dg) plus additional information (Density), are given in this table. The columns marked with a star (*) are estimated values. The G values for 2004 and 2005 are estimated backward from 2009 considering the mean growth between 2006 and 2010 (2011 is a thinning year). The LAIquercus values of LAI-2000 and DHP are calculated using the model proposed by Genet et al. (2010) (Eq. (1)).

| Year | G<br>(m<sup>2</sup>·ha<sup>-1</sup>) |  |  | Age<br>(years) | Density<br>(n·ha<sup>-1</sup>) | Dg*<br>(cm) | LAI<sub>quercus</sub><br>(m<sup>2</sup>·m<sup>-2</sup>) |  |  |  |
| --- | --- | --- | --- | --- | --- | --- | --- | --- | --- | --- |
|  | Q. petraea | C. betulus | Total* |  |  |  | Needles | Litter | DHP* | LAI-2000* |
| 2004 | 20.4 | 3.9 | 24.3 | 120 | 212 | 35.0 | n.d. | n.d. | n.d. | 3.1 |
| 2005 | 20.8 | 4.0 | 24.7 | 121 | 212 | 35.3 | n.d. | n.d. | n.d. | 3.2 |
| 2009 | 22.3 | 4.1 | 26.5 | 125 | 212 | 36.6 | n.d. | n.d. | n.d. | 4.3 |
| 2010 | 22.7 | 4.2 | 26.9 | 126 | 212 | 36.9 | n.d. | n.d. | n.d. | 4.0 |
| 2011 | 19.6 | 3.2 | 22.8 | 127 | 195 | 35.8 | n.d. | n.d. | n.d. | 2.8 |
| 2012 | 19.9 | 3.2 | 23.1 | 128 | 195 | 36.0 | n.d. | 3.4 | n.d. | 2.7 |
| 2013 | 20.2 | 3.3 | 23.5 | 129 | 195 | 36.3 | n.d. | 5.1 | n.d. | 3.5 |
| 2014 | 20.7 | 3.4 | 24.1 | 130 | 195 | 36.8 | n.d. | 4.6 | n.d. | n.d. |
| 2015 | 21.1 | 3.4 | 24.5 | 131 | 195 | 37.1 | n.d. | 4.1 | n.d. | n.d. |
| 2016 | 21.5 | 3.5 | 25.0 | 132 | 195 | 37.5 | 3.0 | 3.5 | n.d. | n.d. |
| 2017 | 21.8 | 3.6 | 25.4 | 133 | 195 | 37.7 | n.d. | 4.1 | n.d. | n.d. |
| 2018 | 22.2 | 3.6 | 25.8 | 134 | 195 | 38.1 | 3.1 | 3.8 | 3.9 | n.d. |
| 2019 | 22.4 | 3.7 | 26.1 | 135 | 195 | 38.3 | n.d. | 3.5 | 3.5 | n.d. |

This site was subject to continuous measurement of the LAI using the litter trap method. From 2012 to 2017, the measurement consisted of 10 collectors of 0.25 m^2^ each, all sorted by species and measured using an LI-3100C Area Meter. Since 2017, the measurement consisted of 20 collectors of 0.5m^2^ each, and all samples were sorted, dried and weighed. Before drying, the contents of the four collectors were measured using an LI-3100C Area Meter. The leaf mass area was the defined and applied to the other collector’s contents to estimate the collected leaf area. The estimated leaf area was then divided by the area covered by all the collectors (10 m^2^) to estimate the stand LAI.

#### Published data from other studies

The data from Le Dantec et al. (2000) used the Li-Cor Plant Canopy Analyzer LAI-2000 detector for four consecutive years (1994-1997) on 17 plots of the Fontainebleau forest (southern Paris region) (Figure 2). Because some of the measurement plots showed mixed stands, the specific LAI of the main species was estimated using Eq. (1)(Genet et al. 2010).

(For details, see Appendix 3.)

The data from Balandier et al. (2006) corresponded to the measurements of canopy transmittance of total solar radiation (TSR) on 30 plots in 5 geographic sites in France (Figure 2). The measurement plots were carefully chosen by the authors on the basis of the (quasi-) absence of concurrent tree species; sufficiently large surfaces and all interference from small understorey trees or herbaceous vegetation were avoided by cutting them off. The measurements were transmittance values, which were converted to LAI using the method described above (see Computation of LAI from measured transmittance).

(For details, see Appendix 4.)

### Models

#### LAI estimation model development

The estimation of the LAI was based on the model developed by (Sonohat et al. 2004). They assumed an age-dependent relationship between LAI and G and age (Eq. (3)):

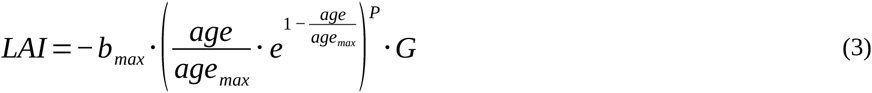

where *LAI* is the estimated LAI, *age* and *G* are the age and basal area, respectively, of the forest stand. The parameters *b*_*max*_ and *P* are fitted against the data, and *age*_*max*_, representing the age at which the LAI is the highest (Balandier et al. 2006), was tested for several values and finally set at 1, for this value gives the best results. This process is detailed in the results section.

Based on this relationship and given the actual LAI/G vs. age relationship observed on our data set, we tested whether a simpler formulation could allow a fair estimation of LAI with fewer parameters (Eq. (4)) :

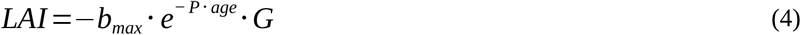

#### Fitting, evaluating and improving the model

All versions of the models were fitted using the nonlinear least squares (NLS) R function, which determines the estimates of the parameters of a non-inear model (the function to minimize is given in Eq. (5)). The evaluation of the model was performed using the “leave-one-out cross-validation” (LOOCV) algorithm in R. The metrics used to evaluate the models were the root mean square error (RMSE), the bias, the modelling efficiency (ME) (Nash and Sutcliffe 1970) and the Akaike information criterion (AIC) (Akaike 1973)

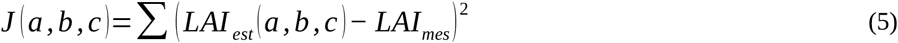

## Results

### Performance of the needle method

The uncertainty of the global measurement of LAI using the needle method decreased rapidly with the first hundred measures. Fifty-two throws were necessary to estimate the LAI with an uncertainty of 0.5 LAI units. To halve this uncertainty, 148 more throws were required to reach 200, and more than 300 throws were required to halve it again. All details of needle measurements is given in Appendix 2.

### Assessment of the LAI-G-age relationship

Equation (6) represents the expression of the age-dependent term. It was chosen by Sonohat et al. (2004) to reflect an age at which the LAI is maximized (age_max_):

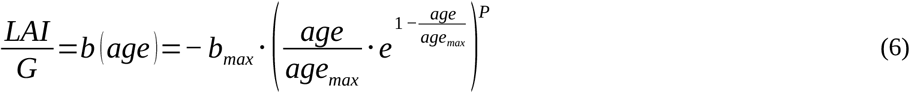

The performance of the best fits given different values of age_max_ showed that 1 is the optimal value of this parameter.

Since age_max_=1 and P(age_max_ = 1) = 0.004 gives the best fit (Figure 4.a, Appendix 5), Eq. (6) can be simplified as a decreasing exponential (Eq. (7)) without significant loss of fit (Figure 4.b, Table 4), as follows:

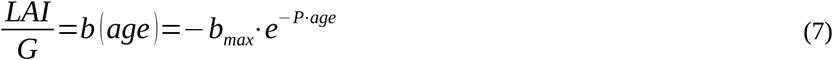

**Figure 4:**
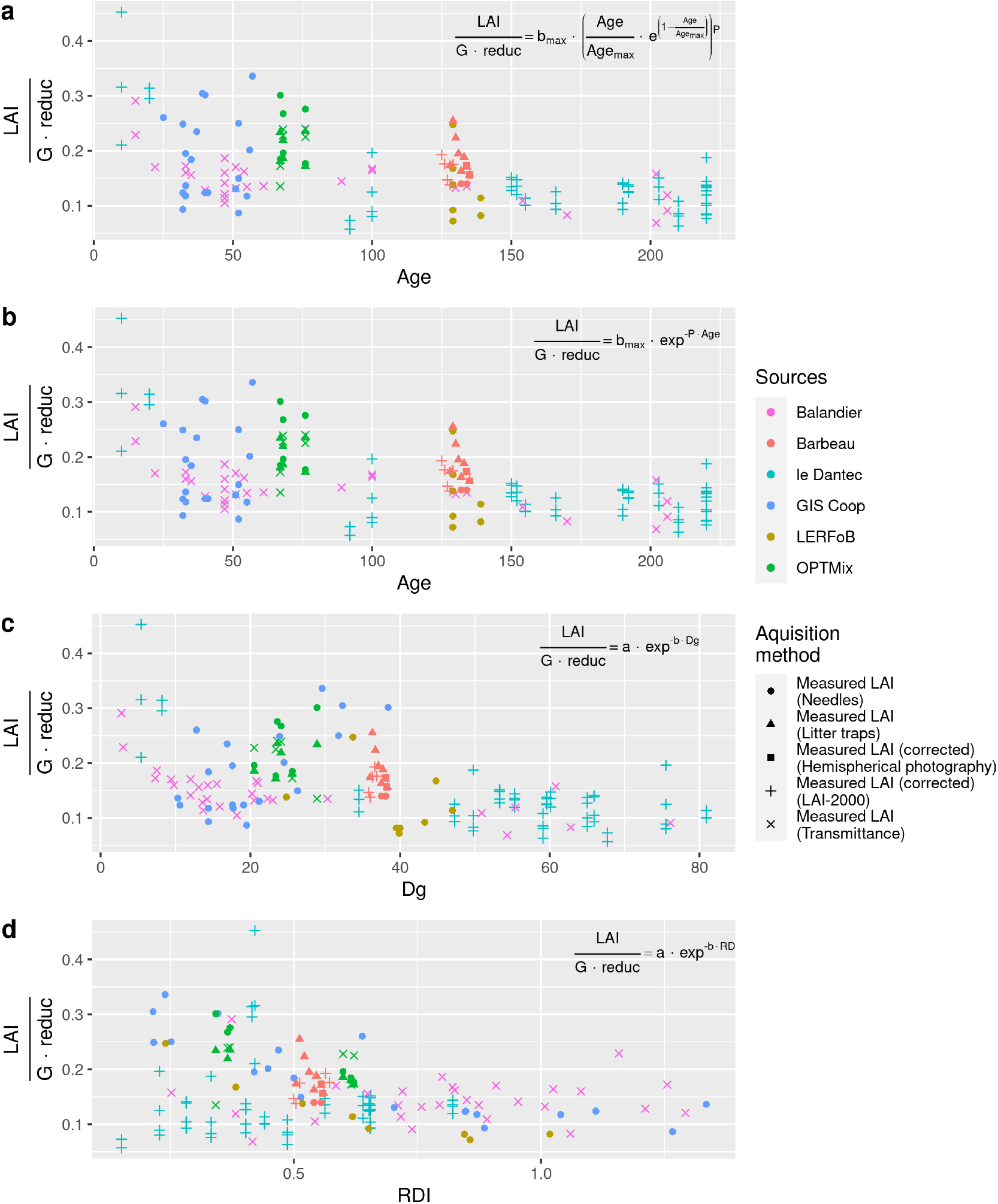
Stand age-(a, b), Dg- (c) and RDI-dependent (d) terms against stand age, Dg and RDI, respectively. The relationship fitted on these figures is displayed in the top-right corner of each panel. The performances are summarized in Table 4. The colours represents the experimental networks and the shapes of the points represents the method used to measure the LAI. Note: the “measured LAI (transmittance)” is the measurement of the transmittance value transformed in LAI value (Eq. (2)) and the DHP and LAI-2000 measured LAI are corrected to the oak-only LAI using the Genet et al. (2010) model (Eq. (1)).

**Table 4:** Statistics of the relationships between LAI:G and age on Dg.

| Relationship | RMSE | Bias | AIC | R<sup>2</sup> |
| --- | --- | --- | --- | --- |
| $b(age) = -b_{max} \cdot \left( \frac{age}{age_{max}} \cdot e^{1 - \frac{age}{age_{max}}} \right)^P$ | 0.05 | 0.00 | -437 | 0.32 |
| $b(age) = -b_{max} \cdot e^{-P \cdot age}$ | 0.05 | 0.00 | -438 | 0.32 |
| $b(Dg) = -a \cdot e^{-b \cdot Dg}$ | 0.05 | 0.00 | -430 | 0.28 |
| $b(RDI) = -a \cdot e^{-b \cdot RDI}$ | 0.06 | 0.00 | -383 | 0.08 |

Replacing the age by the Dg in Eq. (7) did provide a good fit (Table 4 and Figure 4c) but the fit was not as good as the age-dependent relationship. However, even if the relationship between b and Dg appears less parsimonious (higher AIC values, lower R^2^, Table 4), Dg may nevertheless be used accurately when age is not available. The RDI-dependent term showed little to no signal when considering the whole dataset (Table 4 and Figure 4d). Note that the Balandier and Le Dantec data seemed to increase with respect to the RDI, while the GIS Coop, LERFoB and OPTMix data are decreased.

### Assessment and improvement of the model

The estimates of the LAI using the model presented in Eq. (4) are presented in Figure 5. The RMSE was approximately 0.9 points with a nonnull bias of -0.17.

**Figure 5:**
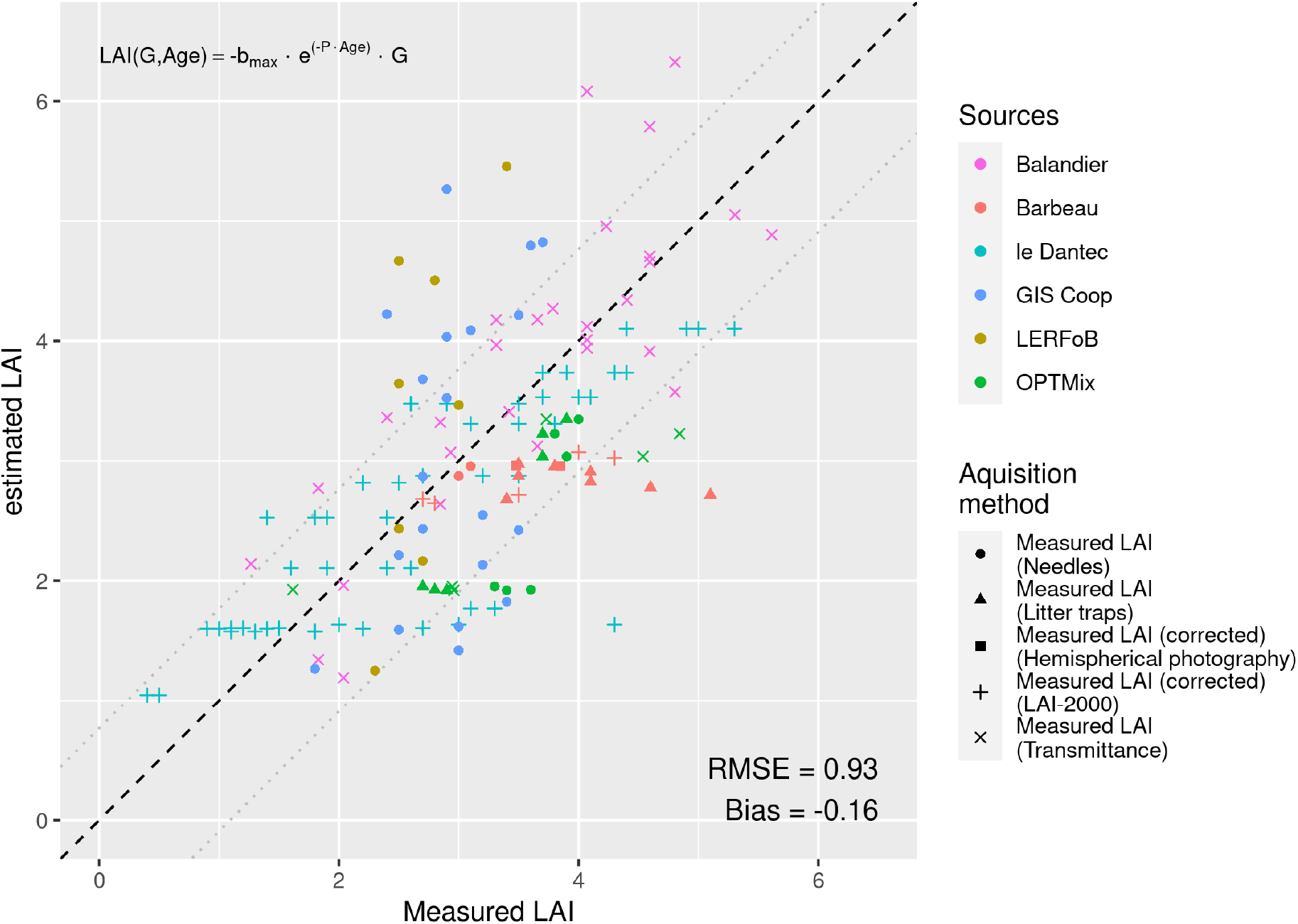
Measured values against estimated ones using the simplified version of the Sonohat et al. (2004) -inspired model (Eq. (4)). The dashed line represent the 1: 1 relation between measured and estimated values and the dotted lines represents the upper (=Bias+RMSE) and lower (=bias-RMSE) values of the confidence range.

The addition of a constant term (Eq. (8)) reduced the RMSE by approximately 14% with a null bias (Figure 6):

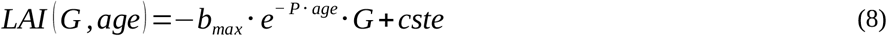

**Figure 6:**
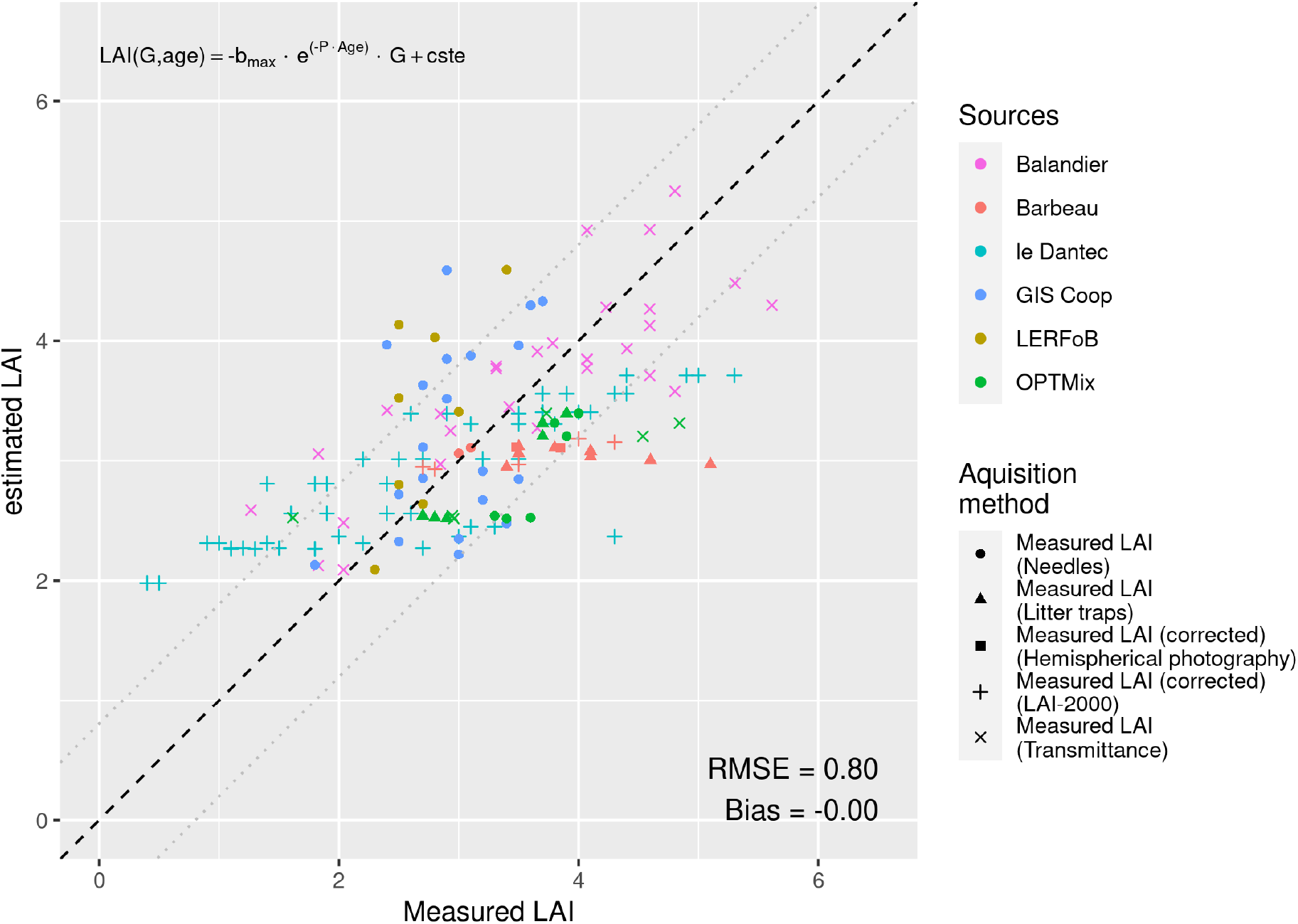
Measured values against estimated ones using the precedent model with addition of a constant term as in Equation (8). The dashed line represent the 1: 1 relationship between measured and estimated values and the dotted lines represents the upper (Bias+RMSE) and lower (bias -RMSE) values of the confidence range.

Replacing the age by Dg (Eq. (9)) gave similar results with an RMSE of 0.77 and a null bias (Figure 7):

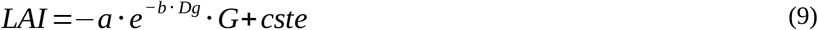

**Figure 7:**
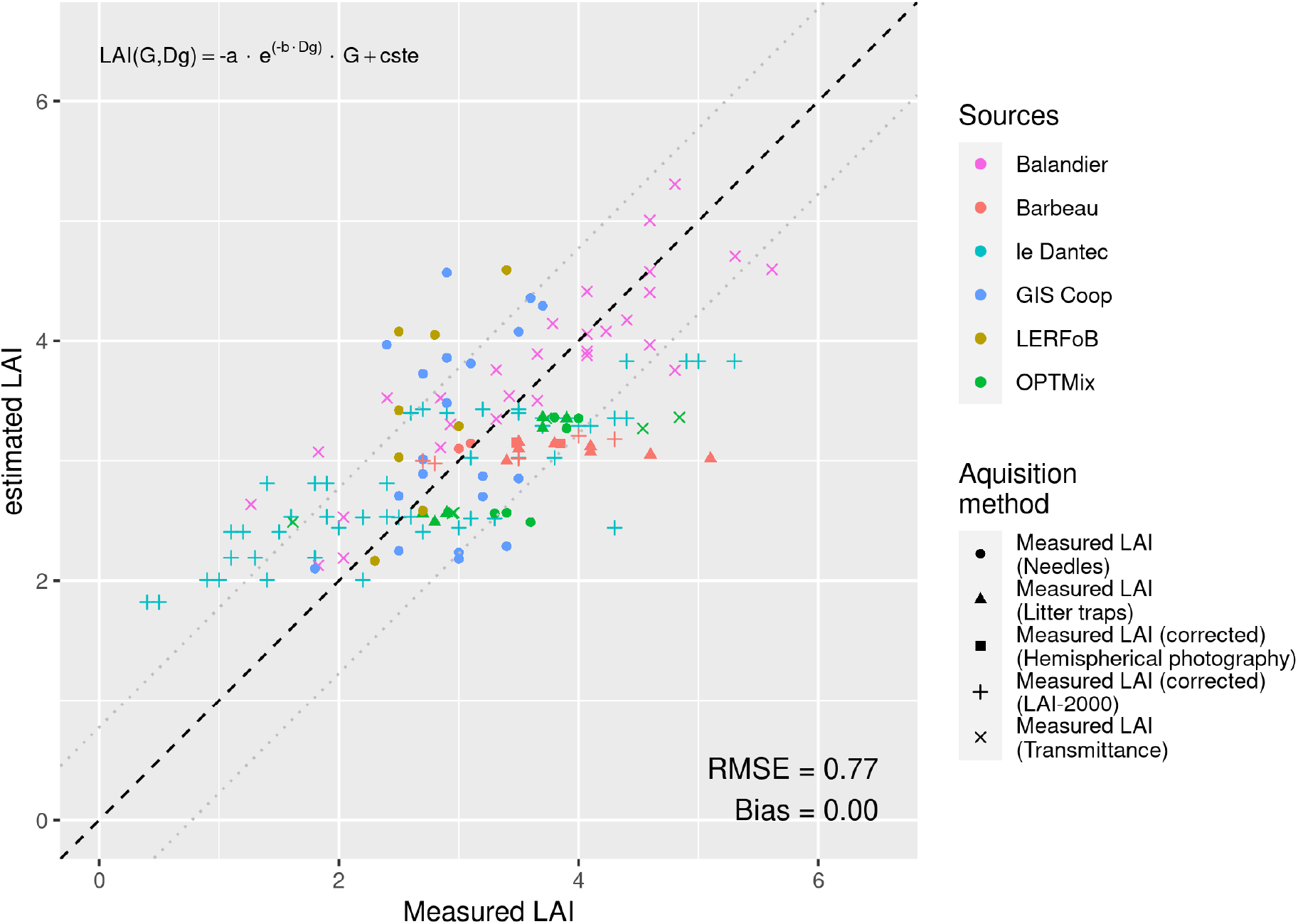
Measured values against estimated ones using the model proposed in Eq. 9. The age-dependent term has been replaced by a Dg-dependant term. The dashed line represent the 1: 1 relationship between measured and estimated values and the dotted lines represents the upper (Bias+RMSE) and lower (bias -RMSE) values of the confidence range.

Note that the distribution of residuals of the model (Eq. (8)) along the Dg range showed a significant linear regression, with a confidence level of 0.95 (Figure 8). The relationship between the residuals and Dg could be included in Eq. (8) to form Eq. (10):

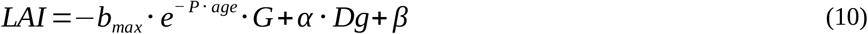

**Figure 8:**
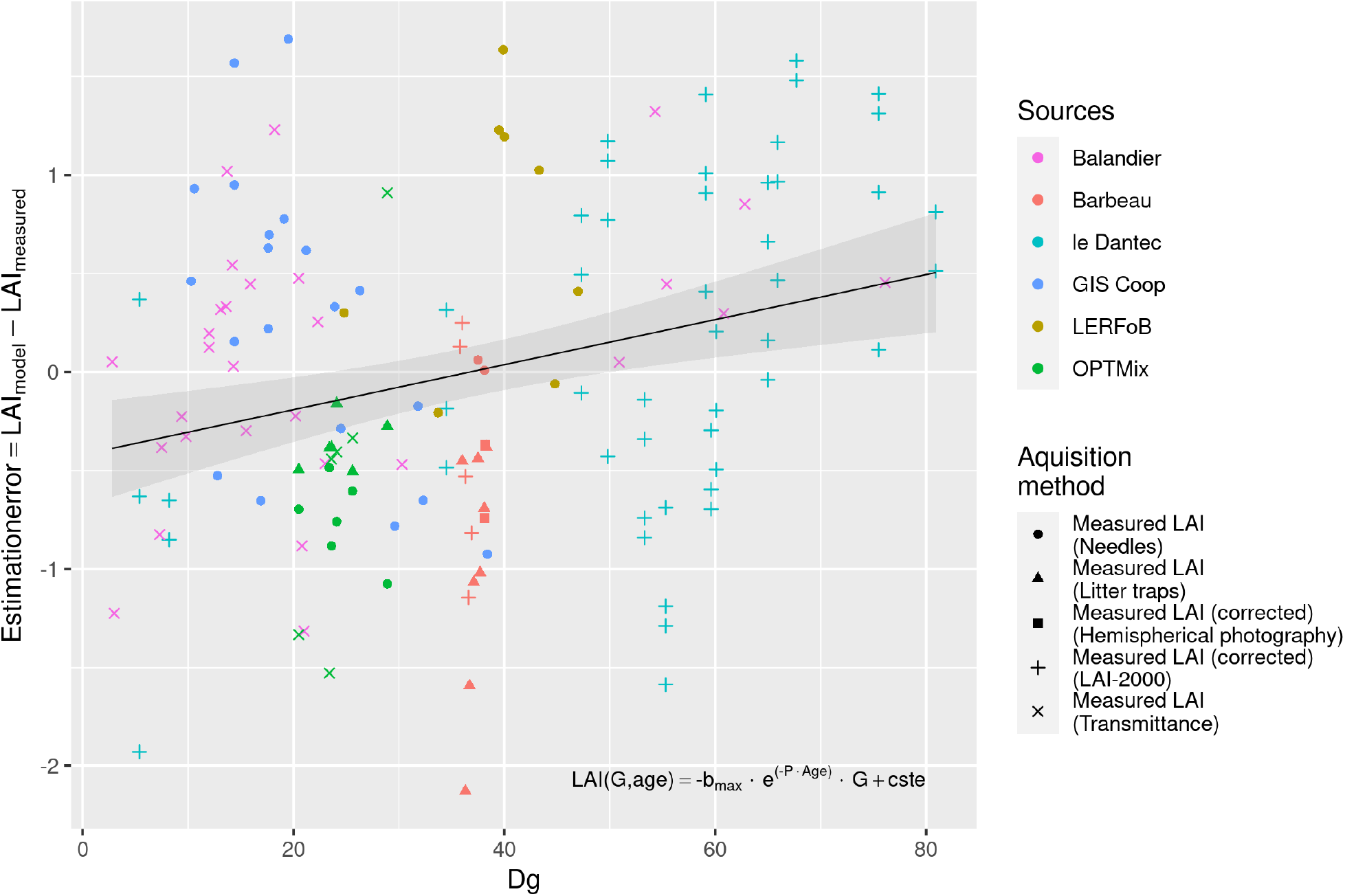
Residuals of the simplified model (Eq. (8)) with constant term against the Dg value. The solid line represents the least-square linear regression and the grey area represents the confidence interval (0.95)

Finally, the Dg-dependent correction factor included in Eq. (10) improved the RMSE by ∼10%. This model fixed the large misestimation of the low-LAI plots measured by Le Dantec et al. (2000)(Figure 9).

**Figure 9:**
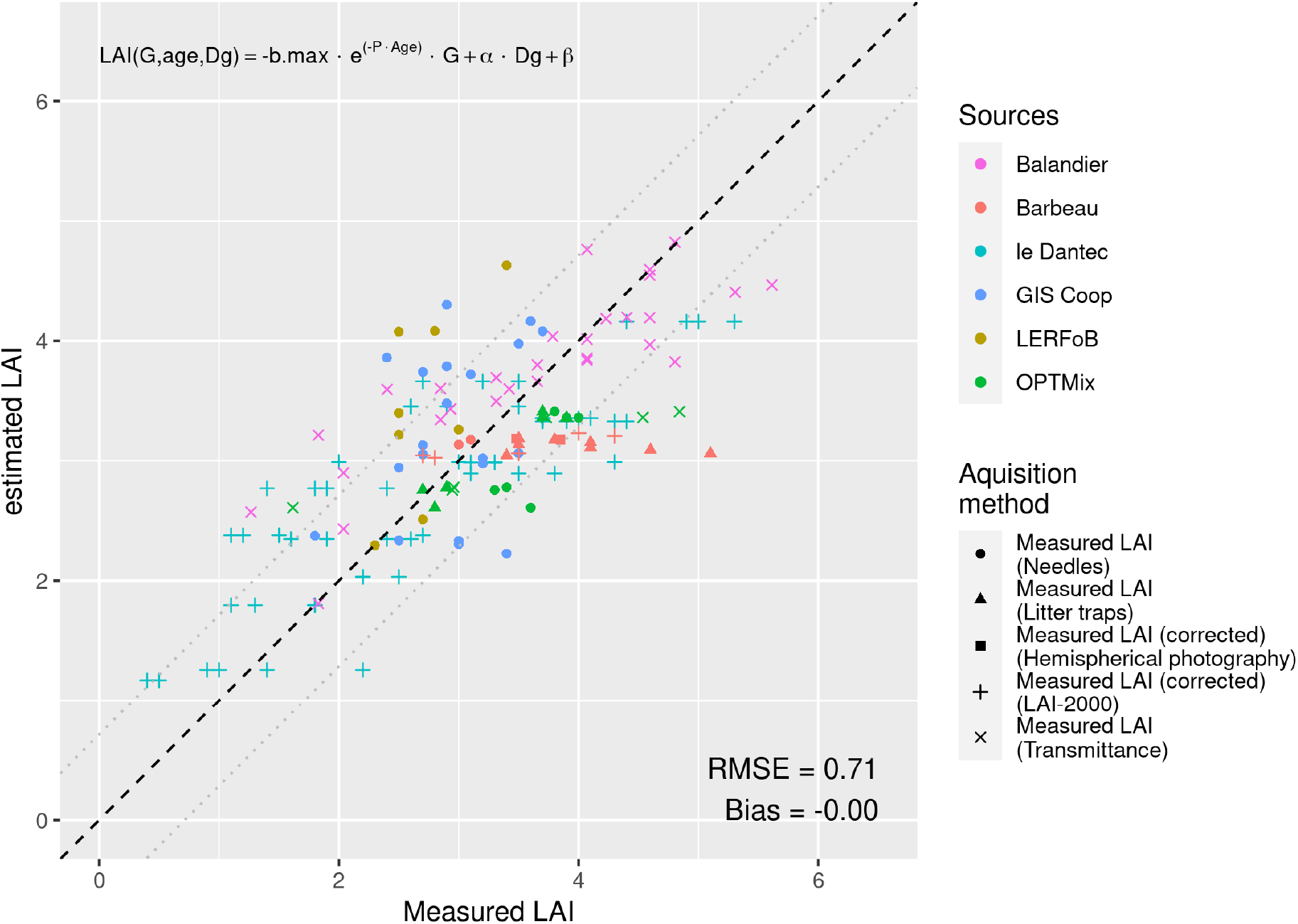
Measured values against estimated ones using the model proposed in Eq. (9). A linear Dg-dependent term is used. The dashed line represent the 1: 1 relation between measured and estimated values and the dotted lines represents the upper (Bias+RMSE) and lower (bias -RMSE) values of the confidence range.

Generally, the use of a constant term was able to cancel the bias in the model.

### LAI measurement methods comparison

The Fontainebleau-Barbeau site has been measured regularly for 10 consecutive years, giving a time series of the evolution of G and Dg (Figure 8). The variables used for our model (G and Dg) did not exhibit the same temporal variations as the measured LAI (in red) (The coefficients of variation for G, Dg and measured LAI were 5%, 2% and 19%, respectively). Therefore, the estimated LAI using our model (black stars, cv = 2%) could not reproduce the temporal variations in the measured LAI.

Overall, note that different measurement methods provided different LAI values except for the DHP that matched the litter trap values for 2018 and 2019 at the Barbeau site. Moreover, the mean LAI over the 2009-2019 period in Barbeau, considering all available measures, was 3.7 (sd = 0.6), while the mean LAI estimation was 3.1 (sd = 0.1). Thus, no significant difference between measures and estimations could be noted under the condition that all measurement methods have the same confidence level.

Figure 11 shows the LAI values measured by different methods in addition to the estimation made by our model described by Eq. (10). When comparing the results obtained by both the litter trap and the LAI-2000 methods, considering the litter trap method as the reference, LAI-2000 tended to underestimate the LAI.

At the OPTMix site, litter trap and needle measurements showed concordant measurements in the high-RDI plots (O12.3, O214.1, O593.1) but diverge in the low RDI plots (O12.2, O214.2, O593.2). Generally, the different measurement methods showed a high degree of variation in their results.

Additionally, our model seemed to underestimate the LAI at both sites (Barbeau and OPTMix) compared to the different measurements (Figure 11).

## Discussion

Based on field and literature data, we proposed an empirical model to estimate the LAI for even-aged oak stands. While there might be a better choice of independent variables (e.g., sapwood area, previous year meteorology) to predict the LAI with greater precision, we chose to propose a method based on widely available and easily obtainable stand variables. This approach allowed the estimation of the stand LAI with fair precision (Table 5). All of the experimental plots measured were even-aged sessile oak stands located in the northern half of France (Figure 2), and since our data span a large range of ages, pedoclimatic conditions and LAI values, nothing seems to forbid the use of this model on a broader area.

**Table 5:**
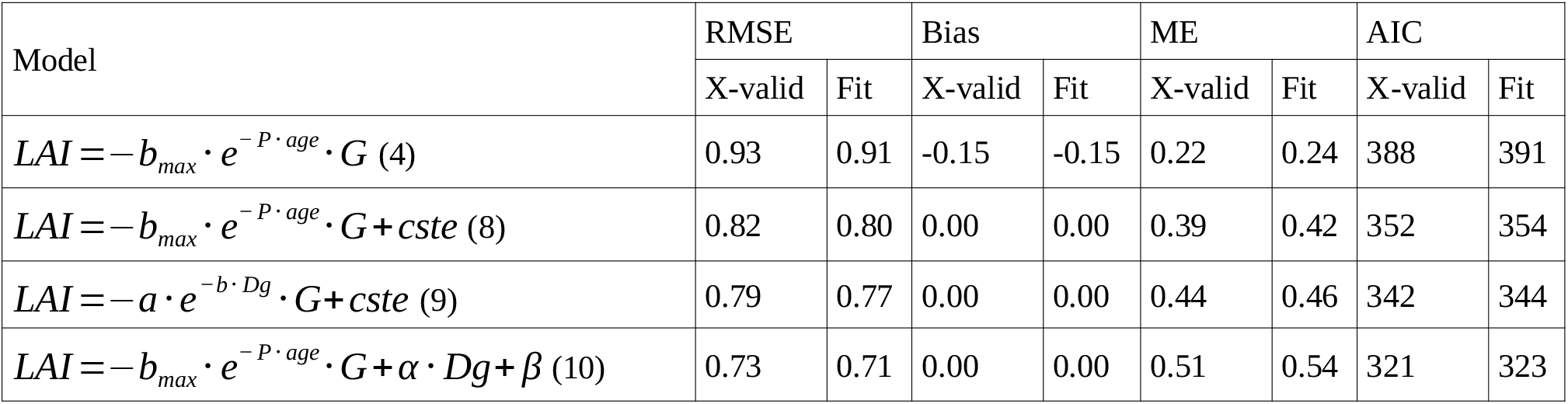
Summary of the statistics of all tested models. Each metric shows its value concerning the cross-validation (X-validation) of the model and the estimation of its parameters (fit). The metrics of each model used in this work are presented in the Appendix 6.

We assumed a relationship between the LAI and the G, Dg and age at the stand level (Jonckheere et al. 2004). Based on this hypothesis, we used the formula developed by Sonohat et al. (2004) on coniferous stands and further used on pure oak stands by Balandier et al. (2006), and we calibrated it to obtain the LAI of pure oak stands. After simplifying it without degrading its predictions, we added a linearly Dg-dependent term and a constant term. In doing so, we managed to cancel its bias and reduce the mean error of its predictions (Table 5).

In forest growth modelling, different methods are used to estimate the LAI. One way to do so is to use net primary production and allow some amount of assimilated carbon to the leaves. The mass of carbon allocated to the canopy is then transformed into leaf dry mass and then leaf area (Running and Gower 1991; Pietsch et al. 2005). Another way is to use the previous year’s LAI and apply a modifier to modulate it given the climatic conditions (condensed in a climatic index relating the soil water stress) (Guillemot et al. 2017). With our method, we proposed estimating a base value of the LAI using a frequently measured stand characteristic. This value could then be adjusted given the environmental conditions.

The Barbeau site measurements showed an interannual variation (measured by both the LAI-2000 from 2009 to 2013 and the litter traps from 2012 to 2019) that was uncorrelated with the evolution of the stand characteristics (Figure 10). By design, this variation enabled the estimation of the mean value of the LAI by averaging the interannual variability of this trait caused by stresses and environmental conditions. If this estimation is made in the context of a process-based model simulation, some modifier should be considered to take into account the perturbation that could happen on such a stand.

**Figure 10:**
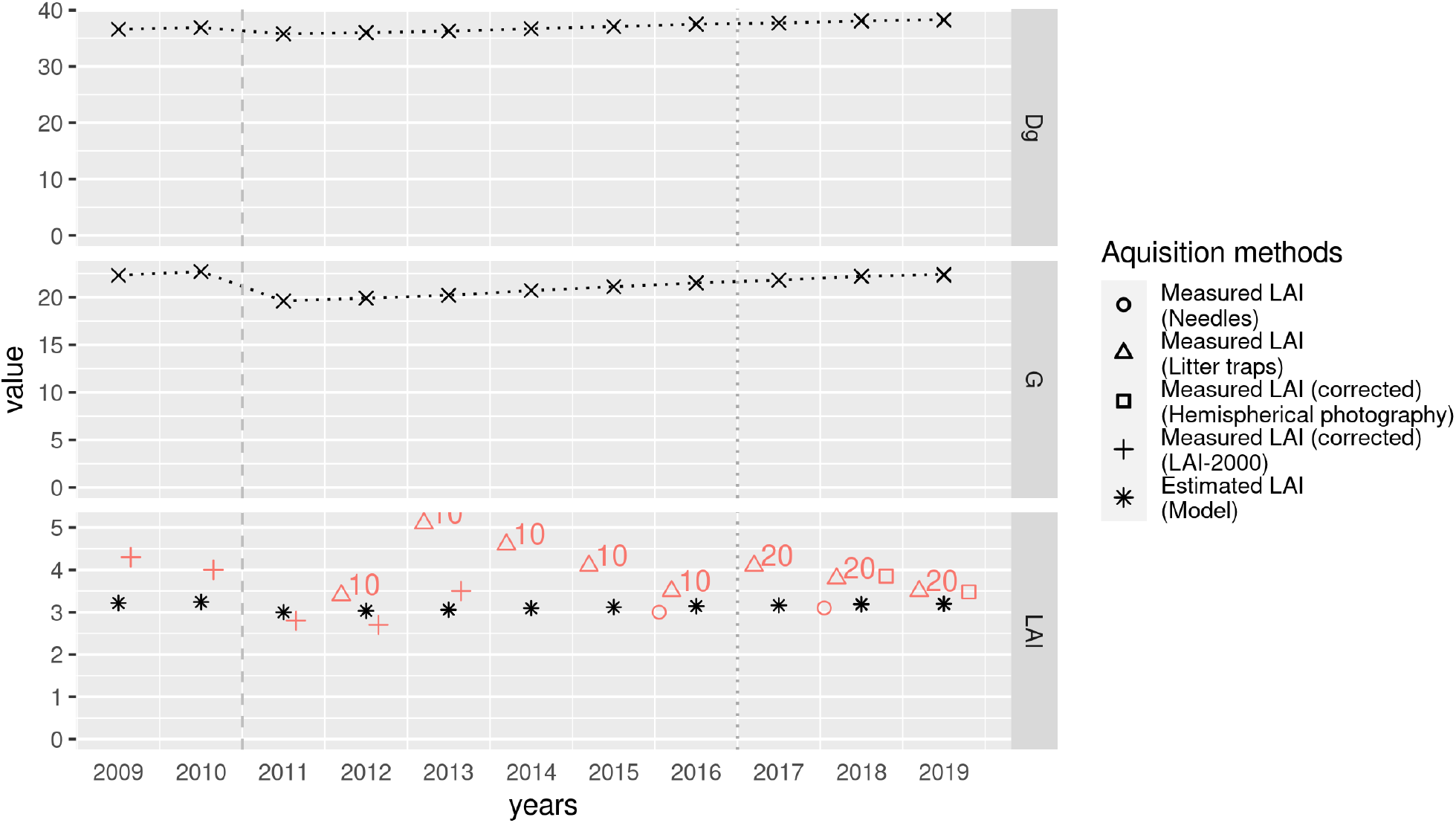
Evolution of the maximum LAI measured on the Barbeau site by different methods (red hollow shapes) against the Dg (top) and G (middle) measured values. The LAI (bottom) estimation based on the G, age and Dg measurement are shown as black stars (*) and are estimated using the model described by Eq. (10). Note the fall of G and measured LAI between the years 2010 and 2011, following a commercial thinning (vertical dashed light-grey line). Also, the litter collection protocol changed slightly (addition of litter traps) during the year 2017.

**Figure 11:**
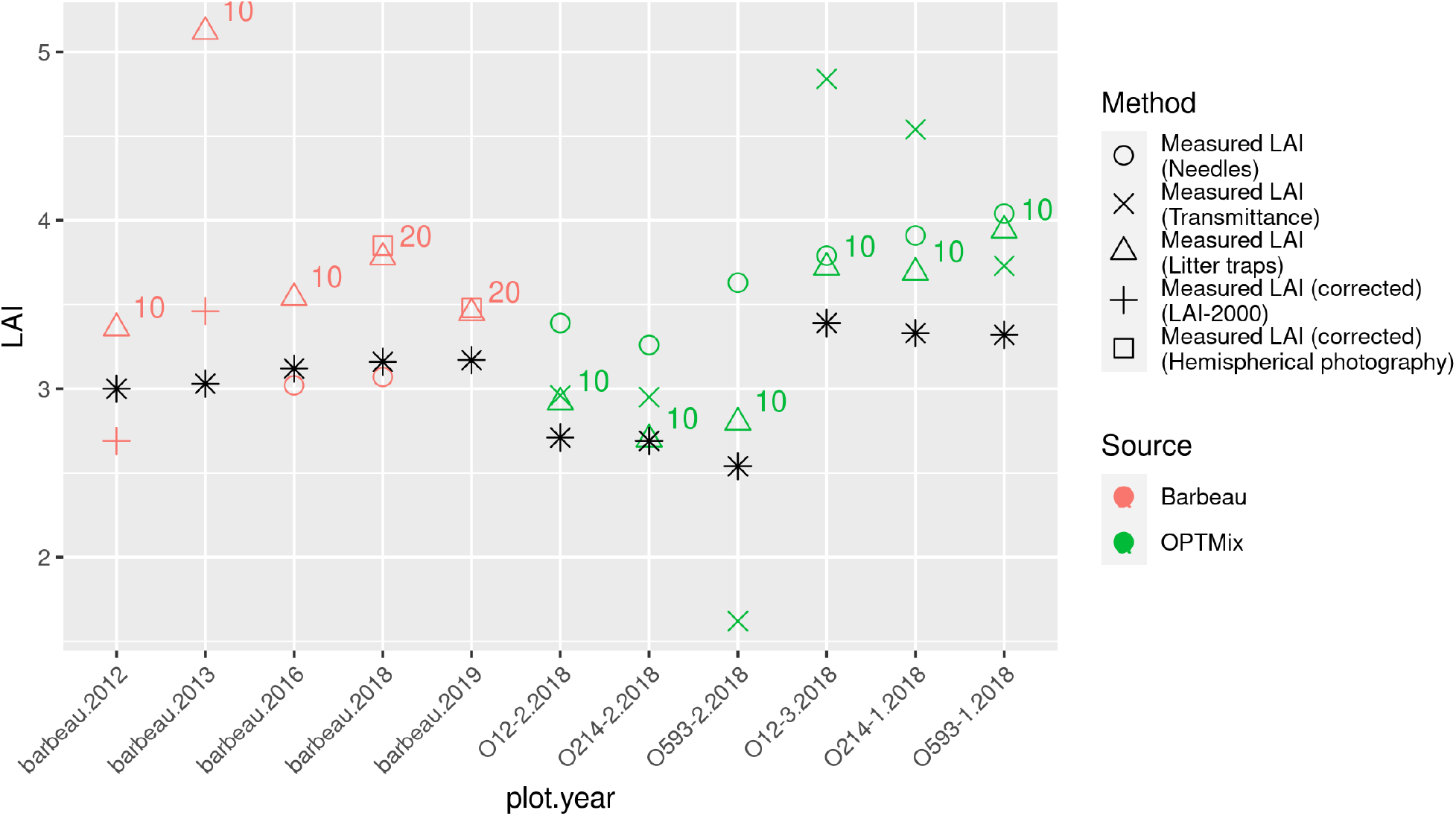
LAI values of Barbeau (one plot, five years) and OPTMix (6 plots, one year) sites using different measurement methods. The shapes represents the methods and the color the site. The black stars (*) are the LAI values estimated using the model described by the Equation (2). Barbeau years presented are only those for which two measures are available to allow comparison between measures as well as between measures and estimation. OPTMix plots are arranged by density with low density first (O12-2, O214-2 and O593-2) and normal density last (O12-3, O214-1 and O593-1).

The needle method, upon which the majority of our original data are based, allows measurement with a fair precision (< 0.5 LAI unit, see Figure 3) of the LAI of a spot at the stand scale. The time needed for two people to collect enough data (about 130 local LAI measurements, see Figure 3) to measure the LAI of one spot is approximately 2 hours of field work. Given the simplicity and genericity of the needed hardware, the needle method is a good method for occasionally measuring the LAI. However, some limitations include the possibility of overestimation by the counting previous year’s leaves as the present year’s leaves in the case of slowly degrading litter and the assumption that the leaves fall homogeneously and exclusively on the stand’s ground. Although the first limitation can be resolved by studying the litter to find the elements allowing distinction between old and new leaves, the second limitation can hardly be resolved other than by defining transects that allow the representation of the heterogeneity of the stand.

Different measurement methods produce different LAI values (Figure 11) for the same plot and year. As the true value of the LAI is unknown and because each measurement method has flaws, it is difficult to assess the LAI measurement methods (Parker 2020). Each is subject to constraints such as a critical sample size (due to the heterogeneity of measurements in the case of litter traps or transmittance measurement) (Bréda 2003; Jonckheere et al. 2004), the time required to process the samples (litter traps) or the hypotheses of spatial and angular distributions of the leaves in the canopy (transmittance measurements, hemispherical photography or LAI-2000) (Weiss et al. 2004).

The comparison between litter traps and LAI-2000 measurements shows the same trend as exposed by Bréda (2003). However, the scarcity of data confirming the underestimation by this method implies caution in this statement.

The use of Dg, G and age prevent the model from predicting interannual variations. There may be many causes of variation, such as pest outbreaks, fire, management thinning operations or climate-related stress (e.g., hight heat, late frost, long drought) (Hogg 1999; Le Dantec et al. 2000; Barr et al. 2004). Additionally, a change in the sampling method, which occurred at the Barbeau site in 2017 (Figure 10), can impact the LAI measurement. That said, the exact nature of the causes of variation is not known, nor is the amplitude each variations would causes. Thus, the model cannot anticipate it given the data used to make predictions.

The present work shows the use of an important set of data using age, density and diameter of trees to predict the LAI of a stand with good precision. Even if the interannual variations for a single stand over a ten-year course are important, the majority of the predictions are close to the measured values (Table 5).

## Acknowledgements

This work was conducted in the context of a doctoral thesis funded by the French National Forest Office (ONF).

This work benefited from the French state aid managed by the ANR under the “Investissements d’avenir” programme with the reference ANR-16-CONV-0003

We thank the ‘GIS Coopérative de données’ (GIS Coop), and specifically the oak group supported by AGROPARISTECH, INRAE and ONF for the management of some of the studied sites and for providing data, and the French Ministry of Agriculture and Forest for GIS Coop financial support.

The OPTMix experimental site where part of our study took place was installed and equipped by the Centre Val-de-Loire region, the Loiret and the French National Forest Office.

The sites belongs to the French national research infrastructure, ANAEE-F (http://www.anaee-france.fr/fr/), and is included in the SOERE TEMPO (https://tempo.pheno.fr/). The sites are also in the framework of the ZAL (LTSER Zone Atelier Loire) and the GIS Coop network (https://www6.inra.fr/GISCoop/), which is supported by the French Ministry for Agriculture and Food.

## Notes

### Competing Interest Statement

The authors have declared no competing interest.

### Summary of Updates

Addition of authors for revision and correction of the aknoledgements to comply with convention.

## References

Akaike H (1973) Maximum likelihood identification of Gaussian autoregressive moving average models. Biometrika 60:255–265

Asner GP, Scurlock JMO, Hicke JA (2003) Global synthesis of leaf area index observations: implications for ecological and remote sensing studies. Global Ecology and Biogeography 12:191–205. https://doi.org/10.1046/j.1466-822X.2003.00026.x

Balandier P, Sonohat G, Sinoquet H, et al (2006) Characterisation, prediction and relationships between different wavebands of solar radiation transmitted in the understorey of even-aged oak (Quercus petraea, Q. robur) stands. Trees 20:363–370. https://doi.org/10.1007/s00468-006-0049-3

Barr AG, Black TA, Hogg EH, et al (2004) Inter-annual variability in the leaf area index of a boreal aspen-hazelnut forest in relation to net ecosystem production. Agricultural and Forest Meteorology 126:237–255. https://doi.org/10.1016/j.agrformet.2004.06.011

Bartelink H (1997) Allometric relationships for biomass and leaf area of beech (Fagus sylvatica L). Annales des Sciences Forestières 54:39–50. https://doi.org/10.1051/forest:19970104

Black TA, Kelliher FM, Wallace JS, et al (1989) Processes controlling understorey evapotranspiration. Philosophical Transactions of the Royal Society of London B, Biological Sciences 324:207–231. https://doi.org/10.1098/rstb.1989.0045

Bréda N, Granier A (1996) Intra-and interannual variations of transpiration, leaf area index and radial growth of a sessile oak stand (Quercus petraea). Ann For Sci 53:521–536. https://doi.org/10.1051/forest:19960232

Bréda N, Soudani K, Bergonzini J-C (2002) Mesure de l’indice foliaire en forêt. GIP ECOFOR, Paris

Bréda NJJ (2003) Ground-based measurements of leaf area index: a review of methods, instruments and current controversies. J Exp Bot 54:2403–2417. https://doi.org/10.1093/jxb/erg263

Bussotti F, Pollastrini M, Holland V, Brüggemann W (2015) Functional traits and adaptive capacity of European forests to climate change. Environmental and Experimental Botany 111:91–113. https://doi.org/10.1016/j.envexpbot.2014.11.006

Cannell MGR, Grace J (2011) Competition for light: detection, measurement, and quantification. Canadian Journal of Forest Research. https://doi.org/10.1139/x93-248

Casa R, Upreti D, Pelosi F (2019) Measurement and estimation of leaf area index (LAI) using commercial instruments and smartphone-based systems. IOP Conference Series: Earth and Environmental Science 275:012006. https://doi.org/10.1088/1755-1315/275/1/012006

Chen JM, Black TA (1992) Defining leaf area index for non-flat leaves. Plant, Cell & Environment 15:421–429

Chen JM, Cihlar J (1995) Plant canopy gap-size analysis theory for improving optical measurements of leaf-area index. Appl Opt, AO 34:6211–6222. https://doi.org/10.1364/AO.34.006211

Clark DB, Olivas PC, Oberbauer SF, et al (2008) First direct landscape-scale measurement of tropical rain forest Leaf Area Index, a key driver of global primary productivity. Ecology Letters 11:163–172. https://doi.org/10.1111/j.1461-0248.2007.01134.x

Cutini A, Matteucci G, Mugnozza GS (1998) Estimation of leaf area index with the Li-Cor LAI 2000 in deciduous forests. Forest Ecology and Management 105:55–65. https://doi.org/10.1016/S0378-1127(97)00269-7

Delpierre N, Berveiller D, Granda E, Dufrêne E (2016) Wood phenology, not carbon input, controls the interannual variability of wood growth in a temperate oak forest. New Phytologist 210:459–470. https://doi.org/10.1111/nph.13771

Dufrêne E, Bréda N (1995) Estimation of deciduous forest leaf area index using direct and indirect methods. Oecologia 104:156–162. https://doi.org/10.1007/BF00328580

Dufrêne E, Davi H, François C, et al (2005) Modelling carbon and water cycles in a beech forest: Part I: Model description and uncertainty analysis on modelled NEE. Ecological Modelling 185:407–436. https://doi.org/10.1016/j.ecolmodel.2005.01.004

Duplat P, Tran-Ha M (1997) Modélisation de la croissance en hauteur dominante du chêne sessile (Quercus petraea Liebl) en France Variabilité inter-régionale et effet de la période récente (1959-1993). Ann For Sci 54:611–634. https://doi.org/10.1051/forest:19970703

Eriksson H, Eklundh L, Hall K, Lindroth A (2005) Estimating LAI in deciduous forest stands. Agricultural and Forest Meteorology 129:27–37. https://doi.org/10.1016/j.agrformet.2004.12.003

Fassnacht KS, Gower ST (1997) Interrelationships among the edaphic and stand characteristics, leaf area index, and aboveground net primary production of upland forest ecosystems in north central Wisconsin. 27:10

Genet H, Bréda N, Dufrêne E (2010) Age-related variation in carbon allocation at tree and stand scales in beech (Fagus sylvatica L.) and sessile oak (Quercus petraea (Matt.) Liebl.) using a chronosequence approach. Tree Physiol 30:177–192. https://doi.org/10.1093/treephys/tpp105

Gower ST, Pongracic S, Landsberg JJ (1996) A global trend in belowground carbon allocation: can we use the relationship at smaller scales? Ecology 77:1750–1755

Granier A, Bréda N, Biron P, Villette S (1999) A lumped water balance model to evaluate duration and intensity of drought constraints in forest stands. Ecological Modelling 116:269–283. https://doi.org/10.1016/S0304-3800(98)00205-1

Guillemot J, Francois C, Hmimina G, et al (2017) Environmental control of carbon allocation matters for modelling forest growth. New Phytologist 214:180–193. https://doi.org/10.1111/nph.14320

Hanewinkel M, Cullmann DA, Schelhaas M-J, et al (2013) Climate change may cause severe loss in the economic value of European forest land. Nature Climate Change 3:203–207. https://doi.org/10.1038/nclimate1687

Hogg EH (1999) Simulation of interannual responses of trembling aspen stands to climatic variation and insect defoliation in western Canada. Ecological Modelling 114:175–193. https://doi.org/10.1016/S0304-3800(98)00150-1

Hurlbert M, Krishnaswamy J, Johnson FX, et al (2019) Risk management and decision making in relation to sustainable development

Jokela EJ, Dougherty PM, Martin TA (2004) Production dynamics of intensively managed loblolly pine stands in the southern United States: a synthesis of seven long-term experiments. Forest Ecology and Management 192:117–130. https://doi.org/10.1016/j.foreco.2004.01.007

Jonckheere I, Fleck S, Nackaerts K, et al (2004) Review of methods for in situ leaf area index determination. Agricultural and Forest Meteorology 121:19–35. https://doi.org/10.1016/j.agrformet.2003.08.027

Korboulewsky N, Pérot T, Balandier P, et al (2015) OPTMix - Dispositif expérimental de suivi à long terme du fonctionnement de la forêt mélangée. 12

Landsberg JJ, Waring RH (1997) A generalised model of forest productivity using simplified concepts of radiation-use efficiency, carbon balance and partitioning. Forest Ecology and Management 95:209–228. https://doi.org/10.1016/S0378-1127(97)00026-1

Le Dantec V, Dufrêne E, Saugier B (2000) Interannual and spatial variation in maximum leaf area index of temperate deciduous stands. Forest Ecology and Management 134:71–81. https://doi.org/10.1016/S0378-1127(99)00246-7

Le Goff N, Ottorini J-M, Ningre F (2011) Evaluation and comparison of size–density relationships for pure even-aged stands of ash (Fraxinus excelsior L.), beech (Fagus silvatica L.), oak (Quercus petraea Liebl.), and sycamore maple (Acer pseudoplatanus L.). Annals of Forest Science 68:461–475. https://doi.org/10.1007/s13595-011-0052-8

Le Moguédec G, Dhôte J-F (2012) Fagacées: a tree-centered growth and yield model for sessile oak (Quercus petraea L.) and common beech (Fagus sylvatica L.). Annals of Forest Science 69:257–269. https://doi.org/10.1007/s13595-011-0157-0

Lindner M, Maroschek M, Netherer S, et al (2010) Climate change impacts, adaptive capacity, and vulnerability of European forest ecosystems. Forest Ecology and Management 259:698–709. https://doi.org/10.1016/j.foreco.2009.09.023

Monserud RA (2003) EVALUATING FOREST MODELS IN A SUSTAINABLE FOREST MANAGEMENT CONTEXT. 1:13

Montagnani L (2018) Ancillary vegetation measurements at ICOS ecosystem stations. International Agrophysics 32:645–664

Morrison IK (2011) Effect of trap dimensions on mass of litterfall collected in an Acersaccharum stand in northern Ontario. Canadian Journal of Forest Research. https://doi.org/10.1139/x91-130

Mussche S, Samson R, Nachtergale L, et al (2001) A comparison of optical and direct methods for monitoring the seasonal dynamics of leaf area index in deciduous forests. Silva Fenn 35:. https://doi.org/10.14214/sf.575

Nash JE, Sutcliffe JV (1970) River flow forecasting through conceptual models part I—A discussion of principles. Journal of hydrology 10:282–290

Ollinger SV (2011) Sources of variability in canopy reflectance and the convergent properties of plants. New Phytologist 189:375–394. https://doi.org/10.1111/j.1469-8137.2010.03536.x

Oswald H (1981) résultats principaux des places d’expérience de chêne du Centre national de Recherches forestières. RFF 33:21

Parker GG (2020) Tamm review: Leaf Area Index (LAI) is both a determinant and a consequence of important processes in vegetation canopies. Forest Ecology and Management 477:118496. https://doi.org/10.1016/j.foreco.2020.118496

Peng C (2000) Understanding the role of forest simulation models in sustainable forest management. Environmental Impact Assessment Review 20:481–501. https://doi.org/10.1016/S0195-9255(99)00044-X

Perot T, Balandier P, Couteau C, et al (2019) Transmitted light as a tool to monitor tree leaf phenology and development applied to Quercus petraea. Agricultural and Forest Meteorology 275:37–46. https://doi.org/10.1016/j.agrformet.2019.05.010

Pietsch SA, Hasenauer H, Thornton PE (2005) BGC-model parameters for tree species growing in central European forests. Forest Ecology and Management 211:264–295. https://doi.org/10.1016/j.foreco.2005.02.046

Reich PB (2012) Key canopy traits drive forest productivity. Proceedings of the Royal Society B: Biological Sciences 279:2128–2134. https://doi.org/10.1098/rspb.2011.2270

Reineke LH (1933) Perfecting a stand-density index for even-aged forests

Running SW, Coughlan JC (1988) A general model of forest ecosystem processes for regional applications I. Hydrologic balance, canopy gas exchange and primary production processes. Ecological Modelling 42:125–154. https://doi.org/10.1016/0304-3800(88)90112-3

Running SW, Gower ST (1991) FOREST-BGC, a general model of forest ecosystem processes for regional applications. II. Dynamic carbon allocation and nitrogen budgets. Tree physiology 9:147–160

Seynave I, Bailly A, Balandier P, et al (2018) GIS Coop: networks of silvicultural trials for supporting forest management under changing environment. Annals of Forest Science 75:48. https://doi.org/10.1007/s13595-018-0692-z

Sindou C, Ruchaud F, Bailly A, et al (2001) Un partenariat scientifique original: la coopérative de données sur la croissance des arbres et peuplements forestiers. Revue forestière française

Skovsgaard JP, Vanclay JK (2008) Forest site productivity: a review of the evolution of dendrometric concepts for even-aged stands. Forestry 81:13–31. https://doi.org/10.1093/forestry/cpm041

Sonohat G, Balandier P, Ruchaud F (2004) Predicting solar radiation transmittance in the understory of even-aged coniferous stands in temperate forests. Annals of Forest Science 61:629–641

Stage AR (1973) Prognosis model for stand development. Intermountain Forest & Range Experiment Station, Forest Service, US …

Verhoef W (1984) Light scattering by leaf layers with application to canopy reflectance modeling: The SAIL model. Remote sensing of environment 16:125–141

Verhoef W (1985) Earth observation modeling based on layer scattering matrices. Remote sensing of environment 17:165–178

Vose JM, Clinton BD, Sullivan NH, Bolstad PV (1995) Vertical leaf area distribution, light transmittance, and application of the Beer–Lambert Law in four mature hardwood stands in the southern Appalachians. Canadian Journal of Forest Research. https://doi.org/10.1139/x95-113

Weiss M, Baret F, Smith GJ, et al (2004) Review of methods for in situ leaf area index (LAI) determination. Agricultural and Forest Meteorology 121:37–53. https://doi.org/10.1016/j.agrformet.2003.08.001

Welles JM (1990) Some indirect methods of estimating canopy structure. Remote sensing reviews 5:31–43

Welles JM, Cohen S (1996) Canopy structure measurement by gap fraction analysis using commercial instrumentation. J Exp Bot 47:1335–1342. https://doi.org/10.1093/jxb/47.9.1335

Wilson JW (1960) Inclined point quadrats. New Phytologist 59:1–7

Yan H, Wang SQ, Billesbach D, et al (2012) Global estimation of evapotranspiration using a leaf area index-based surface energy and water balance model. Remote Sensing of Environment 124:581–595. https://doi.org/10.1016/j.rse.2012.06.004

