## Supplementary material for "Leaf area index estimation of even-aged oak (*Quercus petraea*) forests using in situ stand dendrometric parameters": suplementary data tables

*Appendix 1: Summary of number of LAI data for each source and acquisition method*

| | Balandier | Barbeau | Le Dantec | GIS Coop | LERFoB | OPTMix | Total |
|---|---|---|---|---|---|---|---|
| Needles | - | 2 | - | 20 | 8 | 6 | 36 |
| Litter traps | - | 8 | - | - | - | 6 | 14 |
| Digitial Hemispherical Photography | - | 2 | - | - | - | - | 2 |
| PCA LAI-200 | - | 5 | 53 | - | - | - | 58 |
| Transmittance | 29 | - | - | - | - | - | 29 |
| Total of LAI data | 29 | 17 | 53 | 20 | 8 | 12 | 139 |
| RDI range (min - max) | 0.2 - 1.4 | 0.4 – 0.5 | 0.1 – 0.7 | 0.2 – 1.3 | 0.2 – 0.8 | 0.3 – 0.6 | 0.2 – 1.4 |
| Age range (min, mean, max) | 15 - 206 | 125 - 135 | 10 - 220 | 25 - 57 | 129 - 139 | 67 - 76 | 10 - 220 |
| G range (min, mean, max) | 7 - 49 | 19.6 – 22.7 | 7.2 – 36.8 | 7.7 – 33.4 | 9.3 – 41.4 | 12.6 – 21.9 | 7 – 41.4 |
| Dg range (min, mean, max) | 2.8 – 76.10 | 35.8 – 38.3 | 5.4 – 80.9 | 10.30 – 38.4 | 24.8 – 49.1 | 20.5 – 28.9 | 2.8 – 80.9 |

*Appendix 2: details of the LAI measurements using the needle method. N is the number of local measurements, sd, the estimated error on the LAI and N(sd=0.25) the number of local measurement needed to obtain an estimated error of 0.25 LAI points.*

| Network | Site | plot | N | Average LAI ($m^2. m^{-2}$) | sd | $N_{(sd=0.25)}$ |
|---|---|---|---|---|---|---|
| ICOS | Barbeau | | 120 | 3.1 | 0.34 | 245 |
| GIS Coop | Darney-Sessile | 1 | 126 | 1.8 | 0.30 | 185 |
| GIS Coop | Darney-Sessile | 2 | 128 | 2.5 | 0.35 | 244 |
| GIS Coop | Darney-Sessile | 3 | 133 | 2.9 | 0.40 | 309 |
| GIS Coop | Darney-Sessile | 4 | 132 | 2.5 | 0.35 | 251 |
| GIS Coop | Grosbois | 1 | 117 | 3.5 | 0.42 | 266 |
| GIS Coop | Montrichard | 2 | 149 | 3.5 | 0.38 | 398 |
| GIS Coop | Montrichard | 4 | 133 | 2.7 | 0.35 | 280 |
| GIS Coop | Montrichard | 5 | 129 | 2.7 | 0.43 | 340 |
| GIS Coop | Moulins-Bonsmoulins | 1 | 131 | 3.2 | 0.41 | 374 |
| GIS Coop | Parroy | 1 | 163 | 3.7 | 0.33 | 277 |
| GIS Coop | Parroy | 2 | 141 | 3.1 | 0.33 | 253 |
| GIS Coop | Parroy | 3 | 135 | 3.4 | 0.45 | 386 |
| GIS Coop | Parroy | 4 | 142 | 3.0 | 0.37 | 309 |

| | | | | | | |
|---|---|---|---|---|---|---|
| GIS Coop | Reno-Valdieu | 1 | 126 | 2.7 | 0.43 | 341 |
| GIS Coop | Reno-Valdieu | 2 | 135 | 2.9 | 0.34 | 320 |
| GIS Coop | Reno-Valdieu | 3 | 131 | 2.9 | 0.36 | 290 |
| GIS Coop | Reno-Valdieu | 4 | 131 | 2.5 | 0.35 | 253 |
| GIS Coop | Troncais | 1 | 158 | 3.2 | 0.38 | 329 |
| GIS Coop | Troncais | 2 | 151 | 3.0 | 0.38 | 357 |
| GIS Coop | Troncais | 3 | 161 | 3.6 | 0.33 | 269 |
| OPTMix | O12 | 2 | 131 | 3.4 | 0.46 | 443 |
| OPTMix | O12 | 3 | 124 | 3.8 | 0.42 | 384 |
| OPTMix | O214 | 1 | 120 | 3.9 | 0.42 | 359 |
| OPTMix | O214 | 2 | 120 | 3.3 | 0.48 | 394 |
| OPTMix | O593 | 1 | 126 | 4.0 | 0.44 | 396 |
| OPTMix | O593 | 2 | 122 | 3.6 | 0.51 | 419 |
| LERFoB | Blois | 1 | 155 | 2.7 | 0.28 | 198 |
| LERFoB | Blois | 2A | 124 | 2.3 | 0.39 | 247 |
| LERFoB | Blois | 2B | 131 | 2.5 | 0.31 | 204 |
| LERFoB | Blois | 3 | 145 | 2.5 | 0.30 | 207 |
| LERFoB | Blois | 4 | 143 | 2.5 | 0.27 | 178 |
| LERFoB | Tresor | 1 | 125 | 3.4 | 0.38 | 282 |
| LERFoB | Tresor | 2 | 121 | 3.0 | 0.43 | 260 |
| LERFoB | Tresor | 3 | 122 | 2.8 | 0.40 | 350 |

*Appendix 3: Data from le Dantec (2000). All values necessary for estimating the LAI using our model, measured and estimated (*), are present.*

| Id | $Age_{1994}$ (years) | $G_{total}$ ($m^2 \cdot ha^{-1}$) | $G_{oak}$ ($m^2 \cdot ha^{-1}$) | Density ($N \cdot ha^{-1}$) | Dg * (cm) | $LAI_{total}$ | | | | $LAI_{oak}$* | | | |
|---|---|---|---|---|---|---|---|---|---|---|---|---|---|
| | | | | | | 1994 | 1995 | 1996 | 1997 | 1994 * | 1995 * | 1996 * | 1997 * |
| 1 | 220 | 40.2 | 36.8 | 153 | 55.3 | 6.4 | 6.3 | 6.8 | 5.7 | 5.0 | 4.9 | 5.3 | 4.4 |
| 2 | 150 | 34.2 | 29.0 | 130 | 53.3 | 6.2 | 6.9 | 7.0 | 5.8 | 3.9 | 4.3 | 4.4 | 3.7 |
| 3 | 166 | 35.2 | 27.9 | 159 | 47.3 | 6.6 | 4.9 | 5.4 | 5.0 | 3.5 | 2.6 | 2.9 | 2.6 |
| 4 | 152 | 30.2 | 25.8 | 91 | 60.1 | n. d. | 4.9 | 5.9 | 5.4 | n. d. | 3.1 | 3.8 | 3.5 |
| 5 | 155 | 22.1 | 22.1 | 43 | 80.9 | 2.2 | 2.2 | 2.5 | n. d. | 2.2 | 2.2 | 2.5 | n. d. |
| 6 | 192 | 33.3 | 29.9 | 107 | 59.7 | 5.0 | 5.4 | 5.5 | 5.0 | 3.7 | 4.0 | 4.1 | 3.7 |
| 7 | 210 | 22.2 | 22.2 | 81 | 59.1 | 1.4 | 1.9 | 1.8 | 2.4 | 1.4 | 1.9 | 1.8 | 2.0 |
| 8 | 203 | 25.6 | 24.9 | 266 | 34.5 | n. d. | 3.8 | 3.5 | 2.9 | n. d. | 3.5 | 3.2 | 2.7 |
| 9 | 220 | 18.9 | 18.9 | 57 | 65.0 | 1.9 | 2.6 | 1.6 | 2.4 | 1.9 | 2.6 | 1.6 | 2.4 |
| 10 | 92 | 7.2 | 7.2 | 20 | 67.7 | n. d. | 0.5 | 0.4 | n. d. | n. d. | 0.5 | 0.4 | n. d. |
| 11 | 220 | 15.3 | 14.4 | 74 | 49.8 | 1.4 | 1.4 | 1.8 | 2.7 | 1.2 | 1.1 | 1.5 | 2.7 |
| 12 | 20 | 10.5 | 10.5 | 1988 | 8.2 | n. d. | 3.3 | 3.3 | 3.1 | n. d. | 3.3 | 3.3 | 3.1 |

| | | | | | | | | | | | | | |
|---|---|---|---|---|---|---|---|---|---|---|---|---|---|
| 13 | 10 | 9.5 | 9.5 | 4103 | 5.4 | n. d. | 2.1 | 4.3 | 3.0 | n. d. | 2.0 | 4.3 | 3.0 |
| 16 | 190 | 13.7 | 13.3 | 39 | 65.9 | 1.2 | 1.4 | 2.0 | 1.9 | 1.1 | 1.3 | 1.8 | 1.8 |
| 17 | 100 | 11.7 | 11.2 | 25 | 75.5 | 1.2 | 1.6 | 2.5 | 1.0 | 1.0 | 1.4 | 2.2 | 0.9 |
| 18 | 23 | 25.2 | 25.2 | 5053 | 8.0 | 6.4 | 6.5 | 6.6 | 5.8 | 6.4 | 6.5 | 6.6 | 5.8 |

*Appendix 4: data from Balandier (2006). All values necessary for estimating the LAI using our model, measured and estimated (*), are present.*

| Id | Age (years) | G ($m^2 \cdot ha^{-1}$) | Density ($N \cdot ha^{-1}$) | Dg * (cm) | Transmittance (% incident) | LAI * ($m^2 \cdot m^{-2}$) |
|---|---|---|---|---|---|---|
| 78 | 61 | 27.0 | 694 | 22.3 | 0.16 | 3.7 |
| 86 | 51 | 25.1 | 764 | 20.5 | 0.19 | 3.3 |
| 91 | 51 | 16.7 | 1465 | 12.1 | 0.24 | 2.8 |
| 95 | 134 | 32.6 | 453 | 30.3 | 0.11 | 4.4 |
| 141 | 55 | 21.2 | 1338 | 14.2 | 0.24 | 2.8 |
| 189c | 15 | 7.0 | 11146 | 2.8 | 0.36 | 2.0 |
| 189d | 15 | 21.0 | 30653 | 3.0 | 0.09 | 4.8 |
| 19 | 35 | 18.8 | 1401 | 13.1 | 0.23 | 2.9 |
| 220 | 40 | 35.8 | 2452 | 13.6 | 0.10 | 4.6 |
| 26 | 100 | 34.2 | 991 | 21.0 | 0.06 | 5.6 |
| 47 | 100 | 27.4 | 807 | 20.8 | 0.10 | 4.6 |
| 103c | 47 | 21.4 | 1338 | 14.3 | 0.18 | 3.4 |
| 103d | 47 | 39.7 | 2006 | 15.9 | 0.09 | 4.8 |
| 15 | 154 | 38.8 | 191 | 50.9 | 0.12 | 4.2 |
| 34 | 170 | 49.2 | 159 | 62.8 | 0.13 | 4.1 |
| 36 | 89 | 28.2 | 878 | 20.2 | 0.13 | 4.1 |
| 76 | 54 | 25.1 | 1338 | 15.5 | 0.13 | 4.1 |
| 95 | 22 | 23.9 | 3450 | 9.4 | 0.13 | 4.1 |
| 147c | 33 | 28.7 | 3822 | 9.8 | 0.10 | 4.6 |
| 147d | 33 | 30.8 | 7325 | 7.3 | 0.07 | 5.3 |
| 18 | 206 | 36.4 | 80 | 76.1 | 0.19 | 3.3 |
| 334 | 206 | 17.1 | 71 | 55.4 | 0.36 | 2.0 |
| 347 | 47 | 19.6 | 4459 | 7.5 | 0.16 | 3.7 |
| Colbert | 130 | 34.7 | 835 | 23.0 | 0.10 | 4.6 |
| 7A | 47 | 17.4 | 669 | 18.2 | 0.40 | 1.8 |
| 7B | 47 | 21.1 | 1433 | 13.7 | 0.30 | 2.4 |
| 7C | 47 | 26.8 | 2389 | 12.0 | 0.15 | 3.8 |
| 9A | 202 | 18.5 | 80 | 54.3 | 0.53 | 1.3 |
| 9B | 202 | 11.6 | 40 | 60.8 | 0.40 | 1.8 |

*Appendix 5: Parameters obtained for the different fits on subsets of the available data. The RMSE and bias are calculated for the model prediction versus measurements for the whole set of available data*

| Model | Parameters | Value | Standart error | P-value |
|---|---|---|---|---|
| $LAI = -b_{max} \cdot \left( \frac{age}{age_{max}} \cdot e^{1-\frac{age}{age_{max}}} \right)^{P} \cdot G$ | b. max | 0.1707 | 0.0077 | 2.00E-16 |
| | P | 0.0108 | 0.0021 | 8.87E-07 |
| $LAI = -b_{max} \cdot e^{-P \cdot age} \cdot G$ | b. max | -0.1760 | 0.0088 | 2.00E-16 |
| | P | 0.0021 | 0.0004 | 7.50E-07 |
| $LAI = -b_{max} \cdot e^{-P \cdot age} \cdot G + cste$ | b. max | -0.1093 | 0.0121 | 9.21E-16 |
| | P | 0.0025 | 0.0006 | 4.60E-05 |
| | cste | 1.3758 | 0.1910 | 3.17E-11 |
| $LAI = -a \cdot e^{-b \cdot Dg} \cdot G + cste$ | a | -0.1146 | 0.0112 | 2.00E-16 |
| | b | 0.0102 | 0.0021 | 2.13E-06 |
| | cste | 1.4284 | 0.1799 | 5.61E-13 |
| $LAI = -b_{max} \cdot e^{-P \cdot age} \cdot G + \alpha \cdot Dg + \beta$ | b. max | -0.0643 | 0.0113 | 7.91E-08 |
| | P | -0.0013 | 0.0009 | 1.43E-01 |
| | α | -0.0277 | 0.0045 | 7.55E-09 |
| | β | 2.5390 | 0.2524 | 2.00E-16 |

*Appendix 6: Summary of the metrics of all tested models. "X-valid" columns gives the metrics of the cross validation while the "Fit" gives the metrics of the least-square estimates on all data at once.*

| Model | RMSE | | Bias | | ME | | AIC | |
|---|---|---|---|---|---|---|---|---|
| | X-valid | Fit | X-valid | Fit | X-valid | Fit | X-valid | Fit |
| $LAI = -b_{max} \cdot \left( \frac{age}{age_{max}} \cdot e^{1-\frac{age}{age_{max}}} \right)^{P} \cdot G$ | 0.95 | 0.94 | -0.17 | -0.17 | 0.17 | 0.20 | 397 | 399 |
| $LAI = -b_{max} \cdot e^{-P \cdot age} \cdot G$ | 0.95 | 0.94 | -0.17 | -0.17 | 0.17 | 0.20 | 396 | 399 |
| $LAI = -a \cdot e^{-b \cdot Dg} \cdot G$ | 0.94 | 0.93 | -0.19 | -0.18 | 0.19 | 0.21 | 394 | 396 |
| $LAI = -b_{max} \cdot e^{-P \cdot age} \cdot G + \alpha \cdot Dg$ | 0.95 | 0.93 | -0.14 | -0.13 | 0.18 | 0.21 | 396 | 398 |
| $LAI = -b_{max} \cdot e^{-P \cdot age} \cdot G + cste$ | 0.82 | 0.80 | 0.00 | 0.00 | 0.39 | 0.41 | 354 | 356 |
| $LAI = -a \cdot e^{-b \cdot Dg} \cdot G + cste$ | 0.79 | 0.77 | 0.00 | 0.00 | 0.43 | 0.46 | 343 | 345 |
| $LAI = -b_{max} \cdot e^{-P \cdot age} \cdot G + \alpha \cdot Dg + \beta$ | 0.73 | 0.71 | 0.00 | 0.00 | 0.51 | 0.54 | 321 | 324 |
